# Dissociable contributions of basolateral amygdala and ventrolateral orbitofrontal cortex to flexible learning under uncertainty

**DOI:** 10.1101/2023.04.03.535471

**Authors:** C.G. Aguirre, J.H. Woo, J.L. Romero-Sosa, Z.M. Rivera, A.N. Tejada, J. J. Munier, J. Perez, M. Goldfarb, K. Das, M. Gomez, T. Ye, J. Pannu, K. Evans, P.R. O’Neill, I. Spigelman, A. Soltani, A. Izquierdo

**Author notes:** Co-first authors (C.G. Aguirre, J.H. Woo). **Author contributions:** CGA and AI designed the research; CGA, JLRS, ZMR, ANT, JM, JP, MG, KD, MG, TY, KE, and JP performed the research; CGA, JHW, JM, JLRS, AS and AI analyzed the data; CGA, JLRS, JHW, PRO, IS, AS, and AI interpreted the data; AI, AS, PRO, and IS acquired funding for the project; CGA and AI wrote the paper; all authors edited the final version. **Correspondence**: Claudia Aguirre, Ph.D. or Alicia Izquierdo, Ph.D.

## Abstract

Reversal learning measures the ability to form flexible associations between choice outcomes with stimuli and actions that precede them. This type of learning is thought to rely on several cortical and subcortical areas, including highly interconnected orbitofrontal cortex (OFC) and basolateral amygdala (BLA), and is often impaired in various neuropsychiatric and substance use disorders. However, unique contributions of these regions to stimulus- and action-based reversal learning have not been systematically compared using a chemogenetic approach and particularly before and after the first reversal that introduces new uncertainty. Here, we examined the roles of ventrolateral OFC (vlOFC) and BLA during reversal learning. Male and female rats were prepared with inhibitory DREADDs targeting projection neurons in these regions and tested on a series of deterministic and probabilistic reversals during which they learned about stimulus identity or side (left or right) associated with different reward probabilities. Using a counterbalanced within-subject design, we inhibited these regions prior to reversal sessions. We assessed initial and pre-post reversal changes in performance to measure learning and adjustments to reversals, respectively. We found that inhibition of vlOFC, but not BLA, eliminated adjustments to stimulus-based reversals. Inhibition of BLA, but not vlOFC, selectively impaired action-based probabilistic reversal learning, leaving deterministic reversal learning intact. vlOFC exhibited a sex-dependent role in early adjustment to action-based reversals, but not in overall learning. These results reveal dissociable roles for BLA and vlOFC in flexible learning and highlight a more crucial role for BLA in learning meaningful changes in the reward environment.

**Significance Statement:** Inflexible learning is a feature of several neuropsychiatric disorders. We investigated how the ventrolateral orbitofrontal cortex (vlOFC) and basolateral amygdala (BLA) are involved in learning of stimuli or actions under reinforcement uncertainty. Following chemogenetic inhibition of these regions in both male and females, we measured learning and adjustments to deterministic and probabilistic reversals. For action learning, BLA, but not vlOFC, is needed for probabilistic reversal learning. However, BLA is not necessary for initial probabilistic learning or retention, indicating a critical role for learning of unexpected changes. For stimulus learning, vlOFC, but not BLA, is required for adjustments to reversals, particularly in females. These findings provide insight into the complementary cortico-amygdalar substrates of learning under different forms of uncertainty.

## Introduction

Reversal learning, impacted in various neuropsychiatric conditions, measures subjects’ ability to form flexible associations between stimuli and/or actions with outcomes (Schoenbaum et al., 2003; Izquierdo et al., 2013; Dalton et al., 2016). Reversal learning tasks can also be used to probe learning following expected and unexpected uncertainty in the reward environment (Behrens et al., 2007; Jang et al., 2015; Winstanley and Floresco, 2016; Soltani and Izquierdo, 2019). For example, after the experience of the first reversal, all other reversals are expected to some extent (Jang et al., 2015). Additionally, unexpected uncertainty can be introduced by changes in reward probabilities, after taking the baseline, expected uncertainty into account.

The basolateral amygdala (BLA) is an area of interest in reversal learning due its involvement in value updating (Tye and Janak, 2007; Janak and Tye, 2015; Wassum and Izquierdo, 2015; Groman et al., 2019) and the encoding of both stimulus-outcome and action-outcome associations typically probed in Pavlovian-to-Instrumental tasks (Corbit and Balleine, 2005; Lichtenberg et al., 2017; Malvaez et al., 2019; Sias et al., 2021). Manipulations of amygdala and specifically BLA have resulted in reversal learning impairments (Schoenbaum et al., 2003; Churchwell et al., 2009; Groman et al., 2019), impaired learning from positive feedback (Costa et al., 2016; Groman et al., 2019), enhanced learning from negative feedback (Rudebeck and Murray, 2008; Izquierdo et al., 2013; Taswell et al., 2021), and even improvements of deficits produced by OFC lesions (Stalnaker et al., 2007). Yet BLA has not been extensively studied in the context of flexible reversal learning of stimuli vs. actions with the exception of a recent lesion study in rhesus macaques (Taswell et al., 2021). BLA has also not been systematically evaluated for its contributions to deterministic vs. probabilistic schedules, with the exception of another lesion study in monkeys (Costa et al., 2016). The idea that BLA encodes changes in the environment in terms of salience and associability (Roesch et al., 2010) suggests this region may facilitate rapid updating to incorporate new information. The contribution of BLA to reversal learning and its dependence on the nature of the association (i.e., stimulus-vs. action-based), sensory modality (i.e., visual), and type of uncertainty introduced by the task design (i.e., deterministic vs. probabilistic, but also first reversal versus all subsequent reversals) has also not been extensively studied using a chemogenetic approach.

In parallel, studies with manipulations in rat OFC in reversal learning have included targeting of the entire ventral surface (Izquierdo et al., 2013; Izquierdo, 2017), or more recent systematic comparisons of medial vs. lateral OFC (Hervig et al., 2020; Verharen et al., 2020). Here we examined the role of vlOFC, a subregion not as often probed in reward learning as medial and more (dorso)lateral OFC (cf. Zimmermann et al. (2018)) but also densely-interconnected with BLA (Barreiros et al., 2021a; Barreiros et al., 2021b). Additionally, unlike almost all previous studies on reversal learning, we included both male and female subjects.

Using a within-subject counterbalanced design, we inactivated these regions prior to reversal sessions and measured both learning and adjustments to reversals. We found that vlOFC, but not BLA, inhibition impaired adjustments to deterministic and probabilistic reversals. Conversely, BLA, but not vlOFC, inhibition resulted in significantly slower action-based probabilistic, but not deterministic, reversal learning. Fitting choice data with reinforcement learning models indicated that action-based reversal learning deficits were mediated by a larger memory decay for the unchosen option following vlOFC inhibition, and diminished exploration after reversal following BLA inhibition. These results suggest dissociable roles for BLA in flexible learning under uncertainty, and vlOFC in adjustments to reversals, more generally.

## Materials and Methods

### Subjects

Animals for behavioral experiments were adult (N=70, 33 females; 66 used for behavioral study and 4 males for *ex vivo* imaging) Long-Evans rats (Charles River Laboratories) average age post-natal-day (PND) 65 at the start of experiments, with a 280g body weight minimum for males and 240g body weight minimum for females at the time of surgery and the start of the experiment. Rats were approximately PND 100 (emerging adulthood; Ghasemi et al. (2021)) when behavioral testing commenced. Before any treatment, all rats underwent a 3-day acclimation period during which they were pair-housed and given food and water *ad libitum*. During that time, they remained in their home cage with no experimenter interference. Following this 3-day acclimation period, animals were handled for 10 min per animal for 5 consecutive days. During the handling period, the animals were also provided food and water *ad libitum*. After the handling period, animals were individually-housed under standard housing conditions (room temperature 22–24° C) with a standard 12 h light/dark cycle (lights on at 6am). Animals were then surgerized and tested on discrimination and reversal learning 1-week post-surgery. At the point of reversal, they were beyond the 3-week expression time for Designer Receptors Exclusively Activated by Designer Drugs (DREADDs).

A separate group of Long-Evans rats (N=4, all males) were used for validation of effectiveness of DREADDs in slides of BLA and vlOFC, using *ex vivo* calcium imaging procedures. All procedures were conducted in accordance to the recommendations in the Guide for the Care and Use of Laboratory Animals of the National Institutes of Health and with the approval of the Chancellor’s Animal Research Committee at the University of California, Los Angeles.

### Surgery

#### Viral Constructs

Rats were singly-housed and remained in home cages for 4 weeks prior to testing while the inhibitory hM4Di DREADDs expressed in BLA (n=31, 16 females), vlOFC (n=19, 10 females), or eGFP control virus (n=16, 7 females) in these regions. In rats tested on behavior, an adeno-associated virus AAV8 driving the hM4Di-mCherry sequence under the CaMKIIa promoter was used to express DREADDs bilaterally in BLA neurons (0.1 μl, AP = −2.5; ML= ±5; DV = −7.8 and 0.2 μl, AP= −2.5; ML= ±5; DV= −8.1, from bregma at a rate of 0.1 μl/min; AAV8-CaMKIIa-hM4D(Gi)-mCherry, Addgene, viral prep #50477-AAV8). In other animals, this same virus (AAV8-CaMKIIa-hM4Di-mCherry, Addgene) was bilaterally infused into two sites in vlOFC (0.2 μl, AP = +3.7; ML= ±2.5; DV = −4.6 and 0.15 μl, AP= 4; ML= ±2.5; DV= −4.4, from bregma at a rate of 0.1 μl/min). A virus lacking the hM4Di DREADD gene and only containing the green fluorescent tag eGFP (AAV8-CaMKIIa-eGFP, Addgene) was also infused bilaterally into either BLA (n=7), vlOFC (n=5), or anterior cingulate cortex [(n=5); 0.3 μl, AP = +3.7; ML= ±2.5; DV = −4.6, rate of 0.1 μl/min] as null virus controls. Our vlOFC targeting is most similar to infusion sites reported previously by Dalton et al. (2016) constituting lateral as well as ventral OFC, and 0.7 mm more medial than others (Costa et al., 2023). In rats used for *ex vivo* calcium imaging, the same target regions were infused with either GCaMP6f (AAV9-CaMKIIa-GCaMP6f, Addgene), a 1:1 combination of GCaMP6f+mCherry (AAV8-CamKIIa-mCherry, Vector BioLabs, #VB1947), or a 1:1 combination of GCaMP6f+hM4Di-mCherry (same as used for behavior, AAV8-CaMKIIa-hM4Di-mCherry, Addgene).

#### Surgical Procedure

Infusions of DREADD or eGFP control virus were performed using aseptic stereotaxic techniques under isoflurane gas (1-5% in O_2_) anesthesia prior to any behavioral testing experience. Before surgeries were completed, all animals were administered 5 mg/kg s.c. carprofen (NADA #141–199, Pfizer, Inc., Drug Labeler Code: 000069) and 1cc saline. After being placed in the stereotaxic apparatus (David Kopf; model 306041), the scalp was incised and retracted. The skull was leveled with a +/-0.3 mm tolerance on the A-P to ensure that bregma and lambda were in the same horizontal plane. Small burr holes were drilled in the skull above the infusion target. Virus was bilaterally infused at a rate of 0.01 μl per minute in target regions (coordinates above). After each infusion, 5 min elapsed before exiting the brain.

### Histology

At the end of the experiment, rats were euthanized with an overdose of Euthasol (Euthasol, 0.8 mL, 390 mg/mL pentobarbital, 50 mg/mL phenytoin; Virbac, Fort Worth, TX), were transcardially perfused, and their brains removed for histological processing. Brains were fixed in 10% buffered formalin acetate for 24 h followed by 30% sucrose for 5 days. To visualize hM4Di-mCherry and eGFP expression in BLA or vlOFC cell bodies, free-floating 40-μm coronal sections were mounted onto slides and cover-slipped with mounting medium for DAPI. Slices were visualized using a BZ-X710 microscope (Keyence, Itasca, IL), and analyzed with BZ-X Viewer and analysis software.

Reconstructions of viral expressions of hM4Di (magenta) and green fluorescent protein, eGFP (green) across the AP plane (**Fig. 1BE**) were conducted using Photoshop and Illustrator (Adobe, Inc., San Jose, CA). Two independent raters blind to condition then used ImageJ (U. S. National Institutes of Health, Bethesda, Maryland, USA) to trace and quantify pixels at AP +3.7 (vlOFC) and AP −2.8 (BLA) for each animal. Three measures were obtained per hemisphere and rater measurements were significantly correlated (Pearson correlation: *r* = 0.54, *p* = 4.51e^-04^). There were no differences in expression level between males and females for pixel count reconstructions [*F*_(1,36)_ = 2.59, *p* = 0.12]. Only subjects with bilateral expression were included in behavioral analyses (4 vlOFC and 8 BLA hM4Di rats were excluded due to unilateral expression).

**Figure 1.**
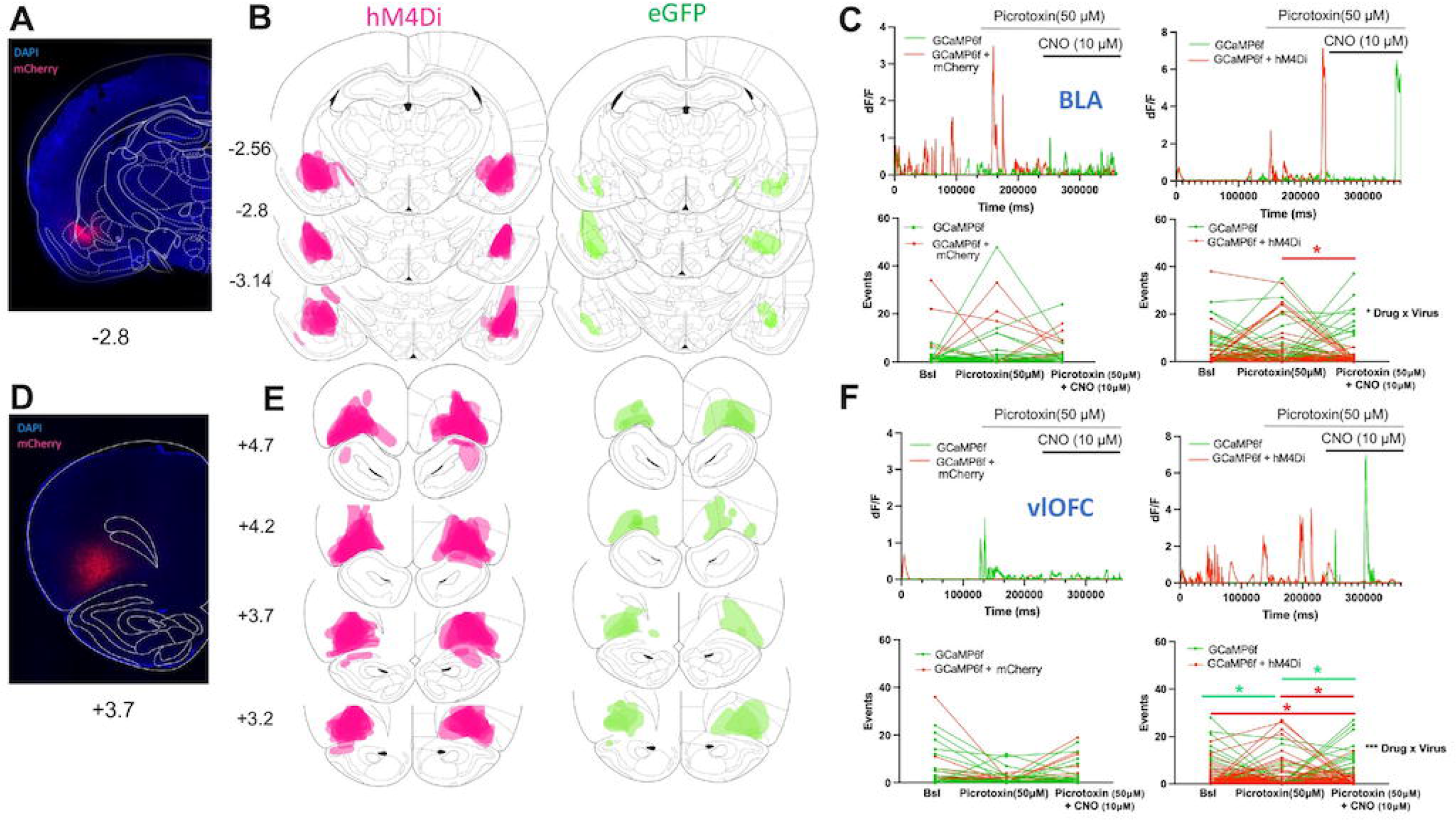
Bilateral targeting of basolateral amygdala (BLA) and ventrolateral orbitofrontal cortex (vlOFC) and confirmation of effective DREADDs inhibition using ex vivo Ca^2+^ imaging in slices. **(A)** Photomicrograph of hM4Di-mCherry DREADDs expression in BLA. Numerals indicate mm anterior to Bregma. **(B)** Reconstructions of viral expression of hM4Di (magenta) and enhanced green fluorescent protein, eGFP (green) in BLA. The more intense colors represent regions where expression overlapped the most across animals. **(C)** In BLA neurons expressing GCaMP6f and GCaMP6f+mCherry, application of CNO (10µM) in the presence of picrotoxin (50 µM) had no effect on the frequency of elicited Ca^2+^ events (top left and right: example of single cell traces, bottom left: Ca^2+^ event changes, each line is a cell). In BLA neurons that expressed hM4Di, there was a reduction in the frequency of elicited Ca^2+^ events during CNO application (bottom right). **(D)** Photomicrograph of hM4Di-mCherry DREADDs expression in vlOFC. Numerals indicate mm anterior to Bregma. **(E)** Reconstruction of viral expression of hM4Di (magenta) and enhanced green fluorescent protein, eGFP (green). The more intense colors represent regions where expression overlapped the most across animals. **(F)** In vlOFC neurons expressing GCaMP6f and GCaMP6f+mCherry, application of CNO (10µM) had no effect on the frequency of elicited Ca^2+^ events (top left and right: example of single cell traces, bottom left: Ca^2+^ event changes, each line is a cell). In vlOFC neurons that expressed hM4Di, there was a reduction in the frequency of Ca^2+^ events during CNO application (bottom right). n=2-5 slices/rat, 2-way ANOVA and multiple comparison tests ***p<0.001, *p<0.05.

### Food Restriction

Five days prior to any behavioral testing, rats were placed on food restriction with females on average maintained on 10-12 grams/ day and males given 12-14 grams/ day of chow. Food restriction level remained unchanged throughout behavioral testing, provided animals completed testing sessions. Water remained freely available in the home cage. Animals were weighed every other day and monitored closely to not fall below 85% of their maximum, free-feeding weight.

### Drug administration

Inhibition of vlOFC or BLA was achieved by systemic administration of clozapine-N-oxide, CNO (i.p., 3mg/kg in 95% saline, 5% DMSO) in animals with DREADDs. Rats with eGFP in these regions underwent identical drug treatment. Rats were randomly assigned to drug treatment groups, irrespective of performance in pretraining. CNO was administered before learning, 30 min prior to behavioral testing. We followed previous work on timing and dose of systemic CNO (Stolyarova, Rakhshan et al., 2019; Hart et al., 2020) and considering the long duration of test sessions. To control for nonspecific effects of injections and handling stress, we also injected animals with saline vehicle (VEH). For reversal learning, to increase power and decrease the number of animals used in experiments, we used a within-subject design for assessing the effects of CNO, with all rats receiving CNO and VEH injections in a counterbalanced order. Thus, for drug administration if a rat received CNO on the first reversal (R1), it was administered VEH on the second reversal (R2), CNO on the third reversal (R3), and VEH on the fourth reversal (R4), or vice versa: VEH on R1, CNO on R2, VEH on R3, and CNO on R4.

### Behavioral Testing

#### Pretraining

Behavioral testing was conducted in operant conditioning chambers outfitted with an LCD touchscreen opposing the sugar pellet dispenser. All chamber equipment was controlled by customized ABET II TOUCH software (Lafayette Instrument Co., Lafayette, IN).

The pretraining protocol, adapted from established procedures (Stolyarova and Izquierdo, 2017), consisted of a series of phases: Habituation, Initiation Touch to Center Training (ITCT), Immediate Reward Training (IMT), designed to train rats to nose poke, initiate a trial, and select a stimulus to obtain a reward (i.e., sucrose pellet). Pretraining stages have been reported in detail elsewhere (Stolyarova, Rakhshan et al., 2019). For habituation pretraining, the criterion for advancement was collection of all 5 sucrose pellets. For ITCT, the criterion to the next stage was set to 60 rewards consumed in 45 min. The criterion for IMT was set to 60 rewards consumed in 45 min across two consecutive days. After completion of all pretraining schedules, rats were advanced to the discrimination (initial) phase of either the action-or stimulus-based reversal learning task, with the task order counterbalanced (**Fig. 2AB** or **Fig. 6AB**). A subset of animals was tested first on the action-based task (13 vlOFC hM4Di, 10 BLA hM4Di), while others were tested on the stimulus-based task first (9 vlOFC hM4Di, 7 BLA hM4Di, 16 eGFP). Three rats in the vlOFC group completed only the stimulus-based task.

**Figure 2.**
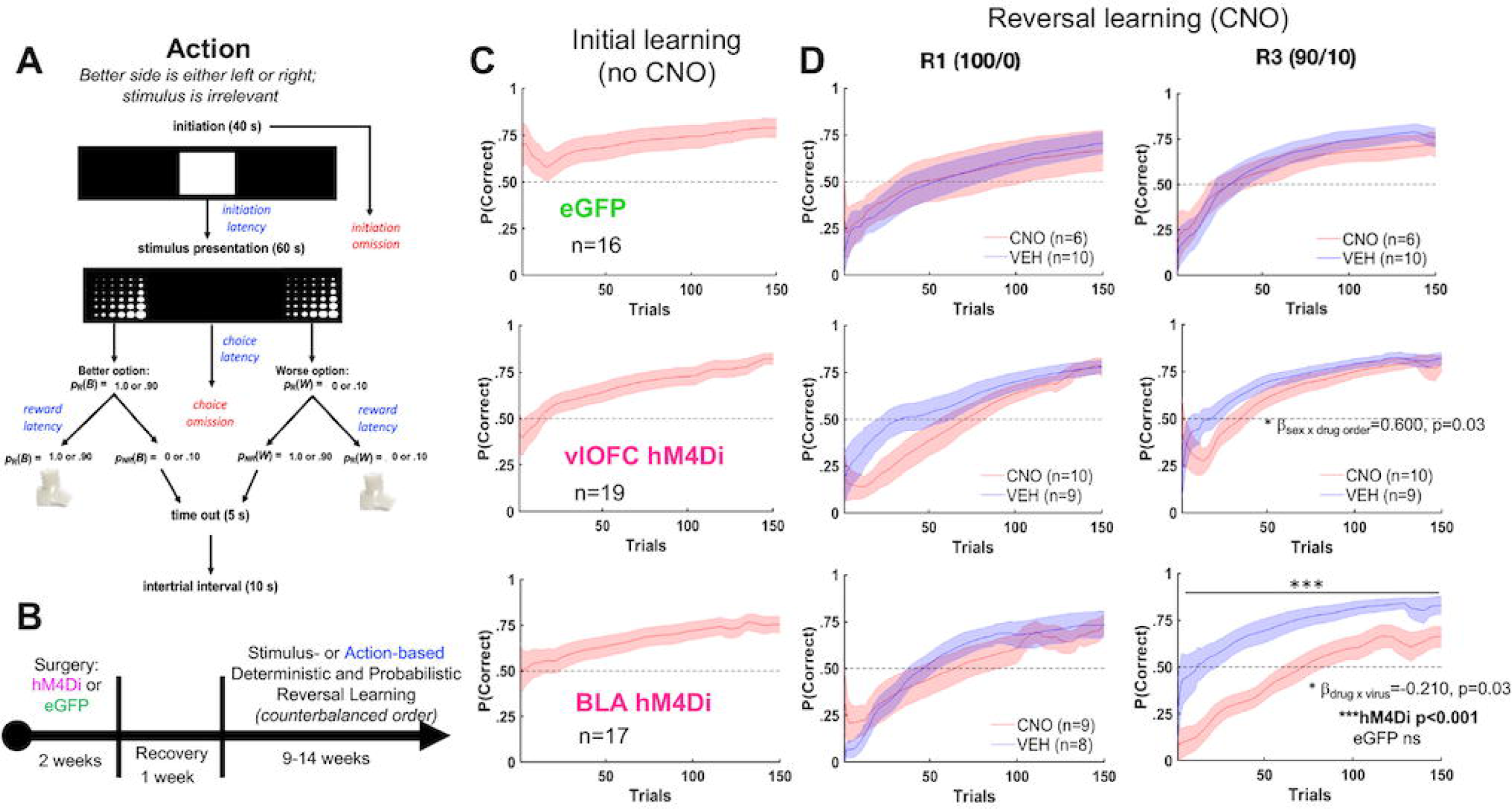
BLA inhibition impairs action-based probabilistic reversal learning, whereas vlOFC inhibition only slows early adjustment to reversals. **(A-B)** Trial structure (A) and timeline (B) of the action-based task. Rats were first surgerized with either hM4Di DREADDs on a CaMKII promoter or eGFP null virus, also on the same promoter. Rats were allowed to recover for 1 week before testing on a stimulus-or action-based reversal learning task. **(C)** Initial learning of a rewarded side. **(D)** Learning during subsequent deterministic (100/0) and probabilistic (90/10) reversal. Plots show cumulative P(Correct) for first 150 trials with a sliding window of 10 trials. Drug order was counterbalanced such that on R2 and R4 animals received VEH if they were administered CNO first on R1 and R3, and vice versa. There was no effect of inhibition on learning in R2 and R4 (not shown). There was no effect of CNO on learning in the eGFP group. *p<0.05 significant sex by drug order interaction (see Fig. 3 for learning curves plotted by sex). Bonferroni-corrected post-hocs following mixed-effects GLM with sex as a covariate fixed factor wherein a drug x virus interaction was found resulted in ***p<0.001 effect of drug only in BLA hM4Di, not in eGFP. The impairment in probabilistic reversal learning following BLA inhibition was also evident in animals with no history of stimulus-based learning and was specific to reversal, not initial probabilistic learning (Fig. 4).

#### Action-based deterministic discrimination learning

After completion of either all pretraining schedules or all four reversals of the stimulus-based task, rats were advanced to the discrimination (initial) phase of the action-based task (**Fig. 2A**). Rats were required to initiate a trial by touching the white graphic stimulus in the center screen (displayed for 40 s), and after initiation rats would be presented with two stimuli (i.e., fan or marble) on the left and right side of the screen (displayed for 60 s). Rats could nosepoke either the spatial side rewarded with one sucrose pellet (the better option, p_R_(B)=1.0;) or the spatial side that went unrewarded (the worse option, p_R_(W)=0.0). Thus, rats were required to ignore the properties of the stimuli and determine the better rewarded side. If a side was not selected, it was scored as a choice omission, and a 10 s inter-trial interval (ITI) ensued. If a trial was not rewarded, a 5 s time-out would occur, followed by a 10 s ITI. If a trial was rewarded, a 10 s ITI would occur after the reward was collected. The criterion was set to 60 or more rewards consumed and selection of the correct option in 75% of the trials or higher during a 60 min session across two consecutive days. After reaching the criterion for the discrimination phase, the rats were advanced to the reversal phase beginning on the next session. Animals were not administered either CNO or VEH injections during discrimination learning.

#### Action-based reversal learning

After the discrimination phase, the rats advanced to the reversal phase. Before a reversal learning session, rats were injected intraperitoneally with either 3 mg/kg of CNO or VEH 30 min prior to each reversal testing session. The side previously associated with the p_R_(B)=1.0 probability was now associated with a p_R_(W)=0.0 probability of being rewarded, and vice versa. The criterion was the same as the deterministic discrimination phase. After reaching the criterion for the first deterministic reversal phase (i.e., R1), the rats advanced to the second deterministic reversal phase (i.e., R2) beginning on the next session. Rats that had previously received VEH during the first reversal would now receive CNO injections, and vice versa.

After completing both deterministic reversal learning phases, rats advanced to the first probabilistic reversal learning phase (i.e., reversal 3, R3). Rats underwent the same injection procedure as the prior reversals. However, the spatial side (i.e., left or right) previously associated with p_R_(B)=1.0, was now associated with a p_R_(W)=0.1 probability of being rewarded, whereas the spatial side previously associated with p_R_(W)=0.0 probability, was now associated with p_R_(B)=0.9. The criterion was the same as the previous deterministic reversal learning phases. After reaching the criterion for the first probabilistic reversal learning phase (i.e., reversal 3, R3), rats were advanced to the second probabilistic reversal phase (i.e., reversal 4, R4) beginning on the next testing day, where the probabilities would be reversed once again. Rats that had previously received VEH during the first probabilistic reversal now received CNO injections, and vice versa. *Stimulus-based deterministic discrimination learning.* After completion of all pretraining schedules (or all reversals of the action-based task), rats were advanced to the discrimination (initial) phase of learning in which they would initiate a trial by touching a white graphic stimulus in the center screen (displayed for 40 s), and choose between two different visual stimuli pseudorandomly presented on the left and right side of the screen, (**Fig. 6A)**. Stimuli were displayed for 60 s each, randomly assigned as the better or worse stimulus: p_R_(B)=1.0 or p_R_(W)=0.0. If a trial was not initiated within 40 s, it was scored as an initiation omission. If a stimulus was not selected, it was scored as a choice omission, and a 10 s ITI ensued. If a trial was not rewarded, a 5 s time-out would occur, followed by a 10 s ITI. Finally, if a trial was rewarded, a 10 s ITI would follow after the reward was collected. A criterion was set to 60 or more rewards consumed and selection of the correct option in 75% of the trials or higher during a 60 min session across two consecutive days. After reaching the criterion for the discrimination phase or if rats were unable to achieve criterion after 10 days, rats were advanced to the reversal phase beginning on the next session. Animals were not administered either CNO or VEH injections during discrimination learning.

#### Stimulus-based reversal learning

After the discrimination phase, rats advanced to the first deterministic reversal learning phase (i.e., reversal 1, R1) where they were required to remap stimulus-reward contingencies. As above, before a reversal learning session, rats were injected intraperitoneally with either 3 mg/kg of CNO or VEH 30 min prior to each reversal testing session. The criterion was the same as discrimination learning. After reaching the criterion for the first reversal phase or if they were unable to achieve criterion after 10 days, the rats were advanced to the second deterministic reversal phase (i.e., reversal 2, R2) beginning on the next testing day, where the reward contingencies were reversed once again. Rats that had previously received VEH during the first reversal now received CNO injections, and vice versa.

After completing both deterministic reversal learning phases, rats advanced to the first probabilistic reversal learning phase (i.e., reversal 3, R3). The injection procedure remained the same as prior reversals. However, the visual stimulus previously associated with p_R_(B)=1.0, would now be associated with p_R_(W)=0.1, whereas the stimulus previously associated with p_R_(W)=0.0, would now be associated with p_R_(B)=0.9. The criterion remained the same as prior reversals. After reaching the criterion for the first probabilistic reversal learning phase or if rats were unable to achieve criterion after 10 days, the rats were advanced to the second probabilistic reversal phase (i.e., reversal 4, R4) beginning on the next testing day, where the probabilities would be reversed once again. As above for action-based reversal learning, rats that had previously received VEH during the first probabilistic reversal now received CNO injections, and vice versa.

#### Action-based probabilistic learning and retention

We assessed initial probabilistic learning, using the same criterion as above, in a separate group of experimentally-naïve animals expressing hM4Di in BLA (N=14, 7 females; 2 animals were excluded due to unilateral expression). Before a learning session, rats were injected intraperitoneally with either 3 mg/kg of CNO (n=10) or VEH (n=2) 30 min prior to testing. One side of the touchscreen was associated with a better probability of reward (p_R_(B)=0.9) and the other side, with a worse probability of reward (p_R_(W)=0.1). Animals were then assessed for their retention of this association on the next session.

### *Ex vivo* calcium imaging

In N=4 animals (all males), following >3 weeks following stereotaxic viral injections, rats (n=1 rat/brain region/virus combination; n=2-5 slices/rat) were deeply anesthetized with isoflurane (Patterson Veterinary, MA, USA), decapitated and brains submerged in ice cold oxygenated (95/5% O_2_/CO_2_) slicing artificial cerebrospinal fluid (ACSF) containing (in mM): 62 NaCl, 3.5 KCl, 1.25 NaH_2_PO_4_, 62 choline chloride, 0.5 CaCl_2_, 3.5 MgCl_2_, 26 NaHCO_3_, 5 N-acetyl L-cysteine, and 5 glucose, pH adjusted to 7.3 with KOH. Acutely microdissected vlOFC or BLA slices (300 μM thick) were obtained (VT1200s, Leica, Buffalo Grove, IL) and transferred to room temperature normal ACSF containing (in mM): 125 NaCl, 2.5 KCl, 1.25 NaH_2_PO_4_, 2 CaCl_2_, 2 MgCl_2_, 26 NaHCO_3_, 10 glucose, pH adjusted to 7.3 with KOH, and allowed equilibrate for >1 hr prior to transfer into a perfusion chamber for imaging.

Imaging was performed on a Scientifica SliceScope, with imaging components built on an Olympus BX51 upright fluorescence microscope equipped with an sCMOS camera (Hamamatsu Orca Flash 4.0v3). Anatomical regions in brain sections for Ca^2+^ imaging were first identified by brightfield imaging with 780nm LED (Scientifica) illumination. Ca^2+^ imaging was performed using a 40x, 0.80NA water immersion objective (Olympus), continuous 470nm LED illumination (ThorLabs), and a filter cube suitable for GCaMP6f imaging: Excitation: Brightline 466/40, Dichroic: Semrock FF495-Di03, Emission: Brightline 525/50. Slices were housed on poly-D lysine cover slips attached to RC-26G chamber (Warner Instruments, Holliston, MA), which was modified with platinum wires to apply electric field stimulation. Images were acquired continually with 20 ms exposure time. Electric field stimulation was applied at 110 mV (twin pulse every 5s). Temperature of ACSF during the recorded sessions was held at 28° C to minimize bubble formation.

### Calcium data extraction

Prior to imaging sessions, 40x images of red and green fluorescence were captured and subsequently overlaid for post-hoc genotyping of individual cells (GCaMP6f^+^, GCaMP6f^+^/hM4Di-mCherry^+^, or GCaMP6f/mCherry^+^). Blinded scorers semi-manually curated regions of interest (ROIs) using Python-based Suite2P software (Pachitariu et al., 2017). ROI fluorescence was subtracted from the annular surround fluorescence, low-pass filtered, and transformed to dF/F_0_ as previously described (Asrican and Song, 2021) where F_0_ is calculated with a boxcar filter with a 200-frame lookback window. dF/F_0_ values were clipped between 0 and 9000 to eliminate negative changes. Area under the curve and event frequency of each cell was calculated for each drug treatment. A threshold of 0.15 dF/F was used to determine significant events, which is lower than the dF/F of a single *ex vivo* action potential, but significantly above signal to noise in our recorded traces (Tada et al., 2014), **Fig. 1CF**.

### Data Analyses

MATLAB (MathWorks, Natick, Massachusetts; Version R2021a) was used for all statistical analyses and figure preparation. Data were analyzed with a series of mixed-effects General Linear Models (GLM, *fitglme*) for the discrimination learning phase to establish there were no baseline differences in learning measures between the hM4Di and eGFP animals (i.e., virus group) prior to any drug treatment for each task separately. Mixed-effects GLMs were also conducted on reversal phases, with all fixed factors included in the model [i.e., reversal number (1-4), virus group (hM4Di, eGFP), drug (CNO, VEH), sex (female, male), drug order (CNO1, VEH1)] and individual rat as a random factor. These GLMs were run for each task type (stimulus- and action-based tasks) separately. Since learning reached asymptote at 5-days for stimulus-based reversal learning, only the first 5 days were included in the GLM. Similarly, since rats typically reached a plateau (and criterion) at 150 trials for action-based reversal learning, we included only the first 150 trials in the GLM. Significant interactions were further analyzed with a narrower set of fixed factors and Bonferroni-corrected post-hoc comparisons. In the instance where sex was found a significant predictor (moderator), sex was entered as a covariate factor in subsequent reversals. Accuracy (probability correct) before and after a reversal (−3 and +3 sessions or −100 trials and +100 trials surrounding a reversal) was analyzed using ANOVA with virus group (hM4Di, eGFP) and drug order (CNO1, VEH1) as fixed factors on the average change pre-post reversal. Virus expression level was analyzed with ANOVA by sex (male, female) on pixel counts obtained via imageJ. Pearson correlation coefficients (*corrcoef*) were analyzed for inter-rater reliability of viral expression quantification. Wilcoxon rank sum tests were used to compare reinforcement learning parameters between groups.

Dependent measures for learning included probability of choosing the correct or better option, probability of win-stay, and probability of lose-switch. Probability of win-stay and lose-switch adaptive strategies were calculated for the stimulus-based task such that each trial was classified as a *win* if an animal received a sucrose pellet, and as a *loss* if no reward was delivered. Statistical significance was noted when p-values were less than 0.05. All Bonferroni post-hoc tests were corrected for number of comparisons.

To analyze *ex vivo* calcium imaging data, 2-way ANOVAs with drug and virus as factors were conducted to compare calcium event changes in GCaMP6f and GCaMP6f+mCherry in each brain region for control experiments, and for GCaMP6f and GCaMP6f+hM4Di-mCherry in each brain region for the experimental group. Tests corrected for number of comparisons were conducted for interactions.

### Reinforcement Learning Models

To capture differences between groups in learning and choice behavior during the action-based task, we utilized two conventional reinforcement learning (RL) models. Specifically, the subjective estimate of reward (*V*) for each choice option was updated on a trial-by-trial basis using reward prediction error (RPE), the discrepancy between actual and expected reward value. In the first model, which we refer to as *RL*, the value estimate of the chosen option (*V_C_*) for a trial *t* was updated using the following equations:

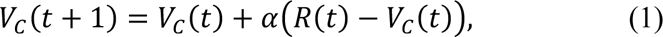

where *R*(*t*) indicates the presence (1) or absence (0) of a reward for the given trial, and α is the learning rate dictating the amount of update in the value estimate by RPE. In this model, the value of unchosen option was not updated.

The second model, referred to as *RL_decay_*, used the same learning rule as Equation (1) for updating the value of the chosen option, and additionally updated the value of the unchosen option (*V_U_*) as follows:

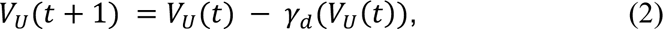

where *γ_d_*is a decay rate controlling the amount of passive decay in value of the unchosen option. In both models above, the probability of choosing a particular option was computed using the following decision rule:

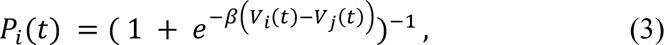

where *i* and *j* corresponds to two alternative options (i.e., left and right for action-based task), and β is the inverse temperature or sensitivity governing the extent to which higher-valued options are consistently selected.

We used the standard maximum likelihood estimation method to fit choice data and estimate the parameters for each session of the experiment. The values of the learning rate α and decay rate *γ_d_*were bounded between 0 and 1, and β was bounded between 1 and 100. Initial parameter values were selected from this range, and fitting was performed using the MATLAB function *fmincon*. For each set of parameters fitted to each session, we repeated 100 different initial conditions selected from evenly-spaced search space to avoid local minima. The best fit was selected from the iteration with the minimum negative log-likelihood (*LL*). For the first model (*RL*), we treated the uninitiated or uncommitted trials with no choice data as if they had not occurred. In contrast, for the second model (*RL_decay_*), both choice options were considered unchosen for those trials and both of the value estimates decayed passively according to Equation (2).

To quantify goodness of fit, we computed both Akaike Information Criterion (AIC) and Bayesian Information Criterion (BIC) for each session as follows:

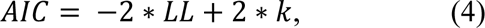

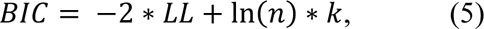

where *k* is the number of parameters in the model (two for *RL* and three for *RL_decay_*), and *n* is the number of choice trials in the session.

## Results

### Ex vivo calcium imaging in slices

We performed ex *vivo* Ca^2+^ imaging to confirm the selective action on CaMKII^+^ neuronal excitability in vlOFC and BLA in rats expressing hM4Di DREADD vs. controls expressing mCherry. In BLA, there was no significant effect of CNO (10µM) on Ca^2+^ events for neurons expressing GCaMP6f or GCaMP6f+mCherry (**Fig. 1C**). A 2-way ANOVA resulted in a significant drug × virus interaction [*F*_(2,324)_ = 3.367, *p* = 0.036], with a selective reduction in the frequency of elicited Ca^2+^ events during CNO only in neurons expressing GCaMP6f+hM4Di (multiple comparison test, *p*=0.049).

In vlOFC, there was also no significant effect of CNO (10µM) on Ca^2+^ events for neurons expressing GCaMP6f or GCaMP6f+mCherry (**Fig. 1F**). However, in CaMKII^+^ vlOFC neurons expressing GCaMP6f+hM4Di there was a decrease in the frequency of Ca^2+^ events during CNO application. A 2-way ANOVA revealed a significant drug x virus interaction [*F* _(2,400)_ = 8.349, *p*< 0.001], with multiple comparisons test resulting in decreased Ca^2+^ events in GCaMP6f+hM4Di following CNO (*p* = 0.02), and increased activity in GCaMP6f expressing neurons after CNO (*p*

= 0.02).

### Discrimination learning: eGFP controls

Mixed-effects GLMs for the discrimination learning phase were conducted for each task separately to establish if there were baseline differences in learning measures between animals infused with eGFP virus in different brain regions. There were no differences between the eGFP groups by target region on learning (i.e., the probability of choosing the correct side) across trials in the action-based task (β_region_ = −0.13, *p* = 0.47), as well as no differences in learning (i.e., probability of choosing the correct visual stimulus) across sessions in the stimulus-based task (β_region_ = 0.10, *p* = 0.09). Thus, animals’ data were collapsed into a single eGFP virus group for subsequent analyses.

### Discrimination learning: hM4Di vs. eGFP

For the action-based task, there were no significant effects of virus or virus interactions for vlOFC vs. eGFP on *probability correct* (β_virus_ = −0.002, *p* = 0.96), with similar findings for the comparison of BLA vs. eGFP (β_virus_ = −0.02, *p*= 0.85; **Fig. 2C**). All animals met criterion very quickly (∼2 days), thus, we compared trials to reach 75% criterion (i.e., probability of choosing the correct side). Both hM4Di virus groups performed comparably [M±SEM: vlOFC hM4Di (69.6±15.4, BLA hM4Di (80.9±21.5)], whereas the eGFP group met criterion within fewer trials (59.5±18.6), but the difference was not statistically significant [vlOFC hM4Di vs. eGFP: β_virus_ = 33.2, *p* = 0.34; BLA hM4Di vs. eGFP: β_virus_ = 51.3, *p* = 0.21].

For the stimulus-based task, there were also no significant effects of virus or virus interactions for either vlOFC vs. eGFP on *probability correct* (β_virus_ = −0.09, *p* = 0.10), or for BLA vs. eGFP (β_virus_ = −0.06, *p* = 0.20; **Fig. 6C**). The animals on average took approximately ∼6 days to meet criterion regardless of virus group [M±SEM: vlOFC hM4Di (6.1±0.7), BLA hM4Di (6.5±1.2), eGFP (6.9±0.7)].

Given the poorer learning in the stimulus-based task, we evaluated whether this was due to the order of task administered [i.e., Stimulus → Action or Action → Stimulus]. To test whether learning was influenced by task order, we analyzed *probability correct* during initial discrimination learning for the stimulus-based task, which resulted in no effect of task order (β_order_ = 0.03, *p* = 0.33), but a significant task order x session interaction (β_order x session_ =-0.03, *p* = 0.002). Thus, subsequent analyses were conducted with task order analyzed separately by session, which revealed that animals administered the Action → Stimulus task order exhibited poorer learning across sessions (β_session_ = 0.01, *p* = 0.05), compared to those administered the Stimulus → Action task order (β_session_ = 0.04, *p* < 0.0001). To further address the possibility that any effects on reversal learning could be due to task order instead of DREADDs inhibition, we conducted a separate assessment of learning in animals administered the Action-based task first (see below).

### Accuracy across trials and sessions: Reversal learning

#### Action-based reversal learning

Mixed-effects GLMs were used to analyze *probability correct,* our primary measure of accuracy, with drug order, virus, and sex as between-subject factors, trial number, reversal number, and drug as within-subject factors, and individual rat as random factor. GLMs were conducted separately by target region (BLA vs. eGFP and vlOFC vs. eGFP), using the following formula for the full model: *γ ∼* [*1 + trial number + reversal number * virus *drug * drug order * sex + (1 + trial number + reversal number * drug| rat)*].

Starting with the full model above for the comparison of BLA with eGFP, we found that trial number was a significant predictor of *probability correct* (β_trial number_ =0.003, *p* = 3.26e^-81^). We also observed a sex x drug order (β_sex x drug order_ =0.63, *p* = 0.01) and sex x virus x drug order (β_sex x virus x drug order_ =-0.72, *p* = 0.046) interaction. To further probe “first reversals” we included only R1 and R3 in the above model. Accordingly, drug order did not vary across R1 and R3. In the analysis of first experiences with deterministic R1 and probabilistic R3, trial number was also a significant predictor of accuracy (β_trial number_ =0.003, *p* = 2.59e^-49^), along with a virus x drug x reversal number interaction (β_virus x drug x reversal number_ =-0.19, *p* = 0.04), which justified further analysis of R1 and R3, separately. Follow-up GLMs were conducted for R1 and R3 *probability correct*, with the following formula: *γ ∼* [*1 + virus *drug + (1 | rat)*]. A significant virus x drug interaction was obtained only for R3: (β_virus x drug_ =-0.22, *p* = 0.03), **Fig. 2D**. Bonferroni-corrected post-hoc comparisons revealed an effect of CNO in hM4Di (*p* = 2.33e^-04^), not in eGFP (*p* = 1.0), with both males and females exhibiting impaired learning of probabilistic R3 following BLA inhibition (**Fig. 3B**). Importantly, stimulus-naïve animals also demonstrated impaired probabilistic reversal learning (β_drug_ =-0.23, *p* = 0.02) (**Fig. 4A**), indicating this slower learning was not due to previous training history.

**Figure 3.**
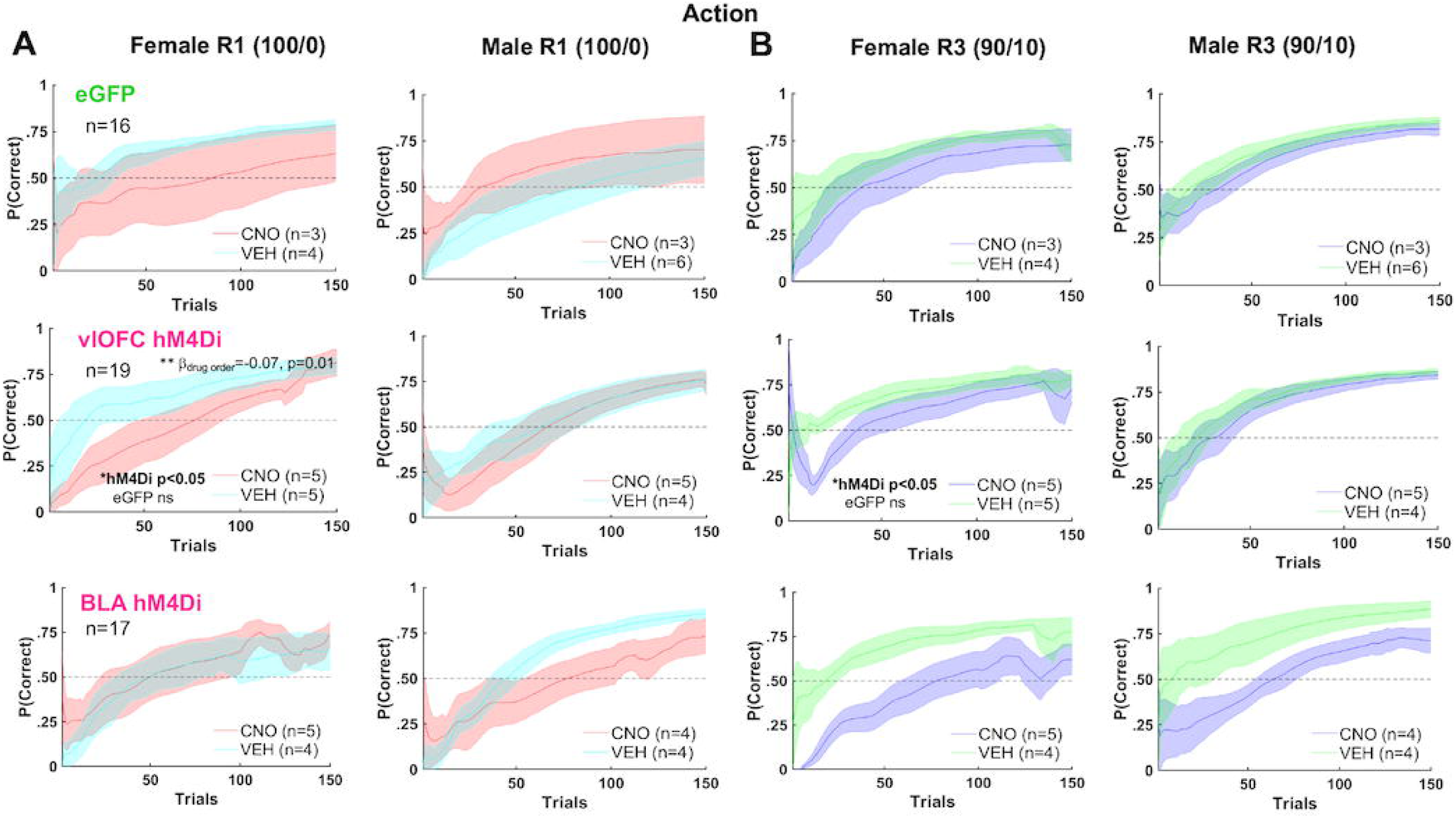
Female adjustment to reversals in the early phase was more affected by vlOFC inhibition than in males, whereas BLA role in probabilistic reversal learning was not sex-dependent. Plotted is the accuracy for first 150 trials with a sliding window of 10 trials. Learning of deterministic (100/0) **(A)** and probabilistic (90/10) **(B)** reversals as measured by probability correct (P(Correct)). Drug order was counterbalanced such that on R2 and R4 (not shown) animals received VEH if they were administered CNO first on R1 and R3, and vice versa. Chemogenetic inhibition of vlOFC lowered P(Correct) in early first deterministic R1 and first probabilistic reversal R3 in females but not males, whereas BLA inhibition attenuated probabilistic reversal learning in both males and females. Bonferroni-corrected post-hocs upon GLM result of sex x drug order interaction resulted in an effect of drug order only in vlOFC females, not in eGFP. The impairing effect of BLA inhibition on learning was not sex-dependent. There were no significant sex differences and no effect of CNO on learning in the eGFP group. **p=0.01, *p<0.05. Reinforcement learning model fit to choice behavior also indicated that chemogenetic inhibition of vlOFC increased the decay rate (decreased memory) of the unchosen option (*γd*) in female rats (Fig. 5).

**Figure 4.**
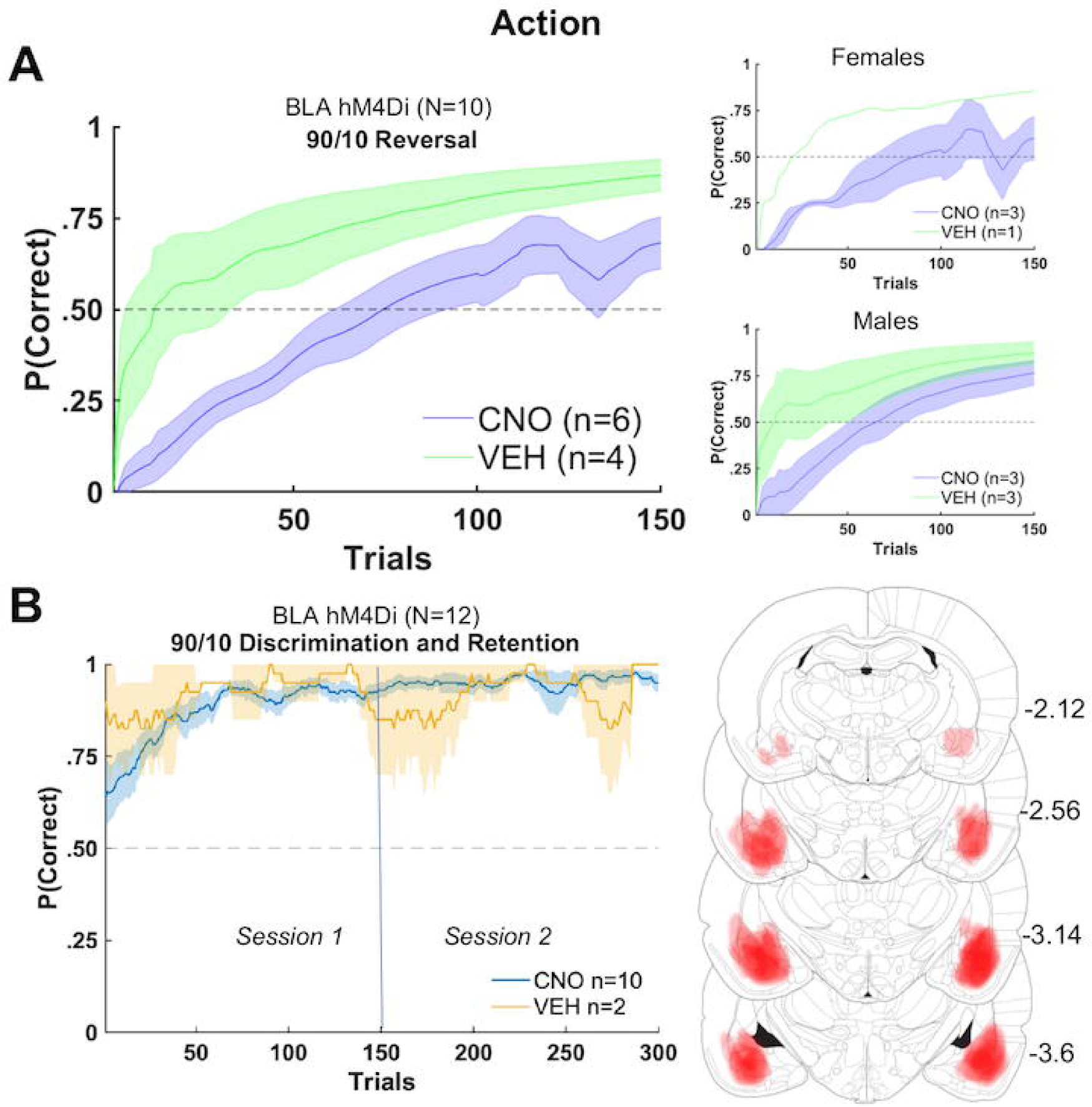
BLA inhibition impairs action-based probabilistic reversal learning in stimulus-naïve animals, but does not affect initial action-based probabilistic discrimination or retention. **(A)** Animals without any prior experience with stimulus-based learning exhibit impaired probabilistic reversal learning. **(B)** There was no significant effect of BLA inhibition on initial probabilistic discrimination or next day retention of the better rewarded side/action. Bilateral expression of hM4Di DREADDs in BLA reconstructed from histology (right) in these animals.

For the comparison of vlOFC with eGFP, trial number was a significant predictor of *probability correct* (β_trial number_ =0.003, *p* = 6.39e^-75^). We also observed a sex x drug order (β_sex x drug order_ =0.60, *p* = 0.03) interaction (**Fig. 2D**). Follow-up GLMs were conducted for accuracy in females and males separately, with the following formula: *γ ∼* [*1 + drug order + (1| rat)*]. A significant drug order effect (CNO first > VEH first) was obtained for females: (β_drug order_ =-0.07, *p* = 0.01), but not males (**Fig. 3**). Due to finding no interactions of reversal number with any other predictor, we were not justified to look further at individual reversals. Altogether, there was no effect of vlOFC inhibition on action-based reversal learning: all animals eventually reached asymptote and criterion over a comparable number of trials.

Since OFC has been implicated in early reversal learning both classically and recently (Jones & Mishkin, 1972; Schoenbaum et al. 2007; Amodeo et al. 2017), we conducted additional analyses using the same full-model formula *γ ∼* [*1 + trial number + reversal number * virus *drug * drug order * sex + (1 + trial number + reversal number * drug| rat)*], but this time restricted to early trials (first 10) of each reversal. Within this early phase, there was a significant sex x virus x drug order (β_sex x virus x drug order_ =-0.94, *p* = 0.03) and sex x virus x drug x drug order interaction (β_sex x virus x drug x drug order_ =1.05, *p* = 0.04). When sex was entered as a covariate, there was a marginally significant interaction of virus x drug order for all reversals (β_virus x drug order_ =-0.22, *p* = 0.056), with a sex difference observed only in the hM4Di group (*p* = 0.03), not in eGFP (*p* = 0.44). Thus, early reversal adjustment in female rats was more adversely affected by previous OFC inhibition (i.e., if they received CNO first), than in males (**Fig. 3**).

#### Action-based probabilistic learning and retention

To further probe if the effect of BLA inhibition on probabilistic reversal learning was specific to reversal, and not initial learning, we administered CNO or VEH before initial 90/10 learning in a separate group of experimentally naïve animals transfected with hM4Di in BLA. To analyze the effect of drug on *probability correct* we used the formula: *γ ∼* [*1 + trial number + drug + (1 + trial number| rat)*]. Only an effect of trial number was found (β_trial number_ =0.001, *p* = 1.7e^-04^). There was no significant effect of drug on learning in either session 1 (initial learning) or session 2 (retention), with or without sex entered in the model as a covariate. Thus, all rats demonstrated learning and full retention on the next day (**Fig. 4B**).

#### Fitting choice behavior in R1 and R3 with Reinforcement learning models

To gain more insight into the effect of vlOFC and BLA inhibition on deterministic reversal learning (R1) and probabilistic reversal learning (R3) and their potential underlying mechanisms, we next compared the estimated model parameters from reinforcement learning models (*RL*, *RL_decay_*). Comparing the goodness of fit between the two models, we found that the second model with the decay parameter (RL_decay_) better accounted for the animals’ choice behavior as indicated by significantly lower AIC (paired sample t-test; t(1334) = 4.613, *p* = 4.34e^-06^). In contrast, the overall mean BIC value was significantly lower for the first model (t(1334) = −2.623, *p* = 8.00 e^-03^). We focused on the estimated parameters from the RL1_decay_ model only (**Fig. 5**), as AIC better distinguished the better-fitting model and allowed estimation of the additional γ*_d_* parameter.

**Figure 5.**
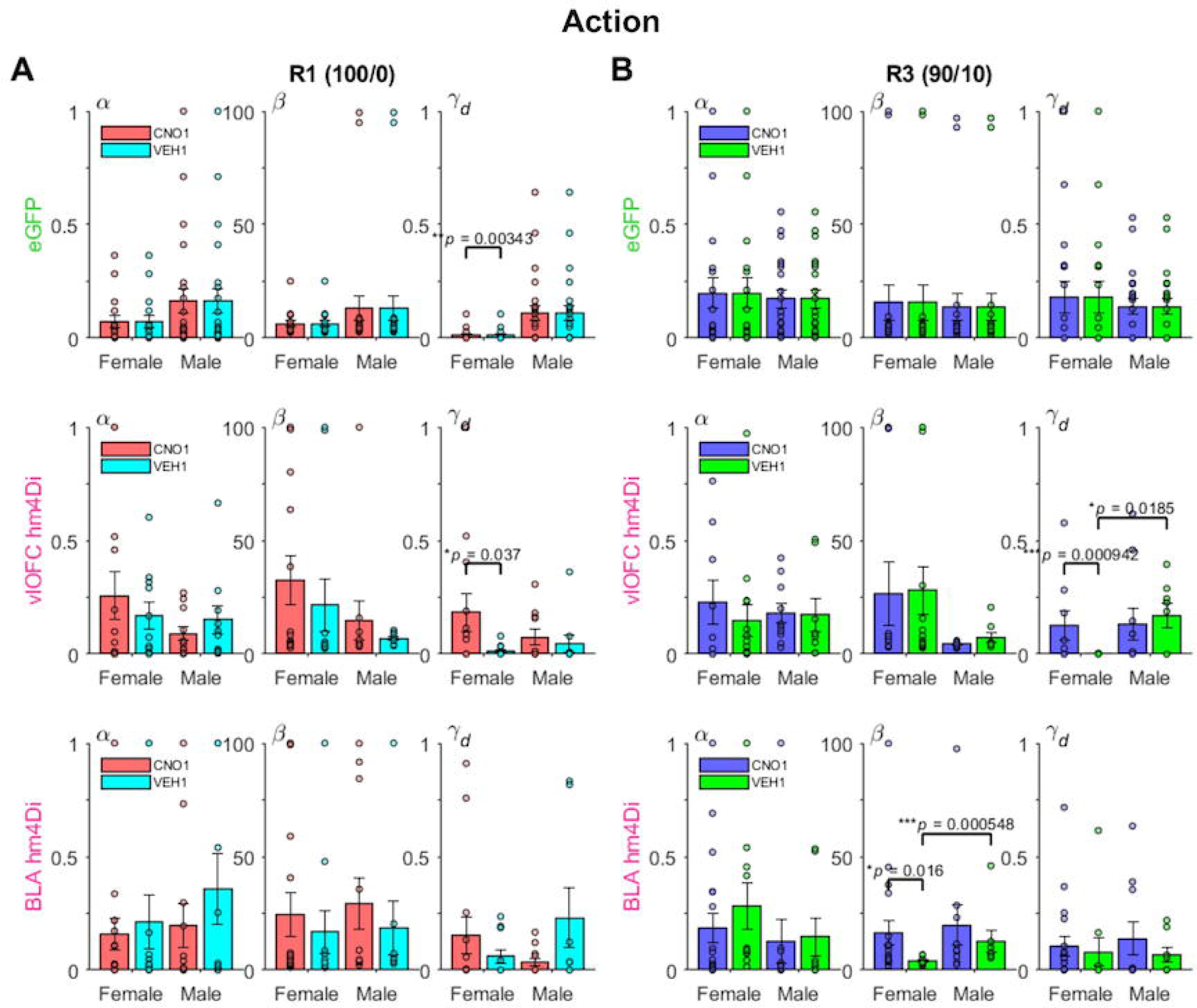
Reinforcement learning model fit to choice behavior indicates differential effects on inverse temperature and decay parameters between male and female rats during deterministic and probabilistic action-based reversals. Plotted are estimated model parameters for RL1decay fitted to each session during deterministic (100/0) **(A)** and probabilistic (90/10) **(B)** reversals. α = learning rate, β = inverse temperature, γd= decay rate for unchosen option. Drug order was counterbalanced such that on R2 and R4 (not shown) animals received VEH if they were administered CNO first on R1 and R3, and vice versa. Female eGFP group exhibited overall lower γd than male eGFP rats during the first deterministic reversal R1. Chemogenetic inhibition of vlOFC increased γd on both reversals (R1 &R3) for female rats, whereas inhibition of BLA increased β on the first probabilistic reversal R3 for female rats. Only comparisons with *p<0.05.

Comparison of the estimated parameters across groups revealed that female and male rats differed mainly in the decay parameter γ*_d_*, which governs the amount of passive decay or forgetting in the value estimate of the unchosen option. During the first deterministic reversal (R1, 100/0), eGFP females showed overall significantly lower values of γ*_d_* than eGFP males (mean difference in γ*_d_* = −0.0982; Wilcoxon rank sum test, *p* = 3.40e^-06^), suggesting a different mechanism of adjustment to the reversal between females and males. While there was no clear evidence for such sex difference in γ*_d_* for the BLA hM4Di groups (mean difference in γ*_d_* = −0.0915; Wilcoxon rank sum test, *p* = 0.24), vlOFC hM4Di groups instead revealed a sex-specific effect between CNO and VEH groups: vlOFC hM4Di females who were administered CNO to inhibit vlOFC had significantly higher values of γ*_d_* compared to those receiving VEH (mean difference in γ*_d_* = 0.170; *p* = 0.04). This effect was even more pronounced during the first probabilistic reversal R3 (mean difference in γ*_d_* = 0.125; *p* = 9.42e^-04^). In contrast, when comparing CNO and VEH in vlOFC hM4Di males there was no difference in γ*_d_* in either R1 (mean difference in γ*_d_* = 0.030; *p* = 0.50) or R3 (mean difference in γ*_d_* = −0.036; *p* = 0.57).

In BLA hM4Di groups, we found a sex-specific effect between CNO and VEH during R3 in the inverse temperature parameter: BLA hM4Di females administered CNO had significantly higher values of β compared to those receiving VEH (mean difference in β = 12.5; *p* = 0.02), indicating higher choice consistency or diminished exploration when the better option had reversed, a maladaptive adjustment to reversal. Yet, BLA hM4Di males did not exhibit this difference (mean difference in β = 7.02; *p* = 0.76). These results based on RL model fitting suggest that the attenuated probabilistic learning in females is mediated by larger β after BLA inhibition and larger γ*_d_* (decreased memory for the unchosen options) after vlOFC inhibition. Importantly, for vlOFC group, this significant difference emerged due to hM4Di VEH females exhibiting enhanced memory in R3.

*Stimulus-based reversal learning.* In contrast to the acquisition curves that demonstrated learning of the initial visual discrimination (**Fig. 6C**), all animals exhibited difficulty with stimulus-based reversal learning, rarely achieving above 60% after 10 sessions (**Fig. 7**), similar to recent reports (Harris, Aguirre et al., 2021; Ye et al., 2023). Here, due to several non-learners, we adhered to the criterion of rats reaching greater than a 50% running window average for the last 100 trials in discrimination, for inclusion in subsequent reversal learning analyses. The following numbers did not meet this criterion and were excluded from these groups: 0 of 10 vlOFC hM4Di, 3 of 10 BLA hM4Di, and 5 of 16 eGFP. GLMs were conducted separately in only these ‘stimulus learners’ for accuracy (*probability correct*) by target region comparison, but with session instead of trial number as a within-subject factor. Thus, the GLM formula for the full model was as follows: *γ ∼ [1 + session *reversal number * virus * drug * drug order * sex + (1 + session * reversal number * drug| rat)*].

**Figure 6.**
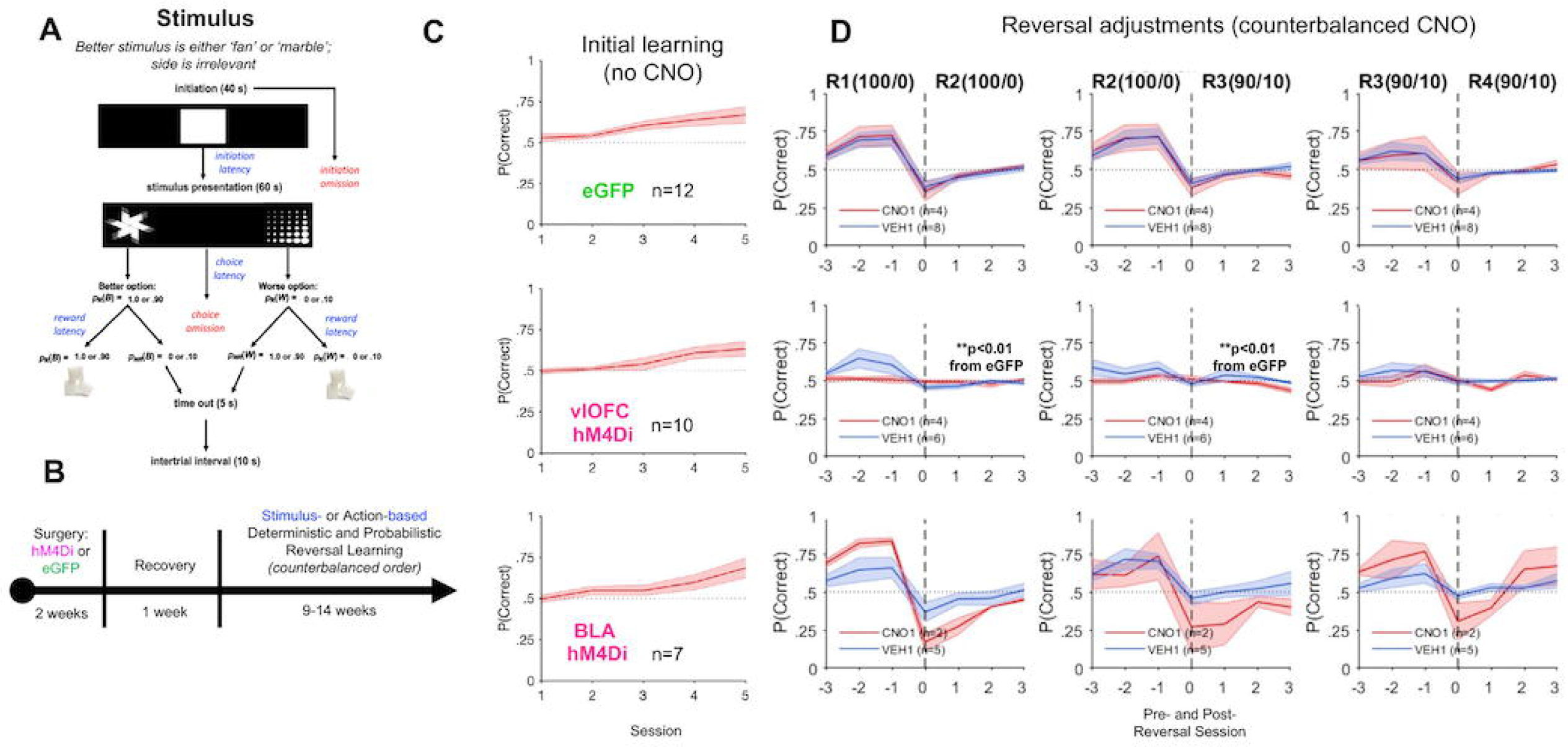
vlOFC, but not BLA, inhibition impairs adjustment to stimulus-based reversals as measured by probability correct. (**A-B**) Trial structure (A) and timeline (B) of the stimulus-based task. Rats were first surgerized with either hM4Di DREADDs or eGFP null virus on a CaMKII promoter. Rats were allowed to recover for 1 week before testing on a stimulus-or action-based reversal learning task. (**C**) Initial learning of a rewarded stimulus, presented pseudorandomly on the left or right side of the touchscreen for eGFP (top), vlOFC hM4Di (middle), and BLA hM4Di (bottom). (**D**) Plots of accuracy around reversals were restricted to include only animals that reached greater than 50% running window average for the last 100 trials in initial discrimination. Rats were always tested on a deterministic schedule before a probabilistic one. Shown are subsequent deterministic (100/0) and probabilistic (90/10) reversal “transitions,” 3 sessions before and after each reversal. Drug order was counterbalanced such that on R2 and R4 animals received VEH if they were administered CNO first on R1 and R3, and vice versa. Chemogenetic inhibition of vlOFC abolishes changes in probability correct over the last 3 (pre) and first 3 (post)-reversal sessions, indicating impaired adjustments to reversals. In contrast, BLA inhibition had no clear impact on reversal learning. There was also no effect of CNO in eGFP group learning. **p<0.01 different than eGFP following ANOVA of pre-post difference. vlOFC inhibition abolished learning after the first stimulus-based reversal (Fig. 7), but did not significantly impair adjustment around action-based probabilistic reversal as following BLA inhibition (Fig. 8).

**Figure 7.**
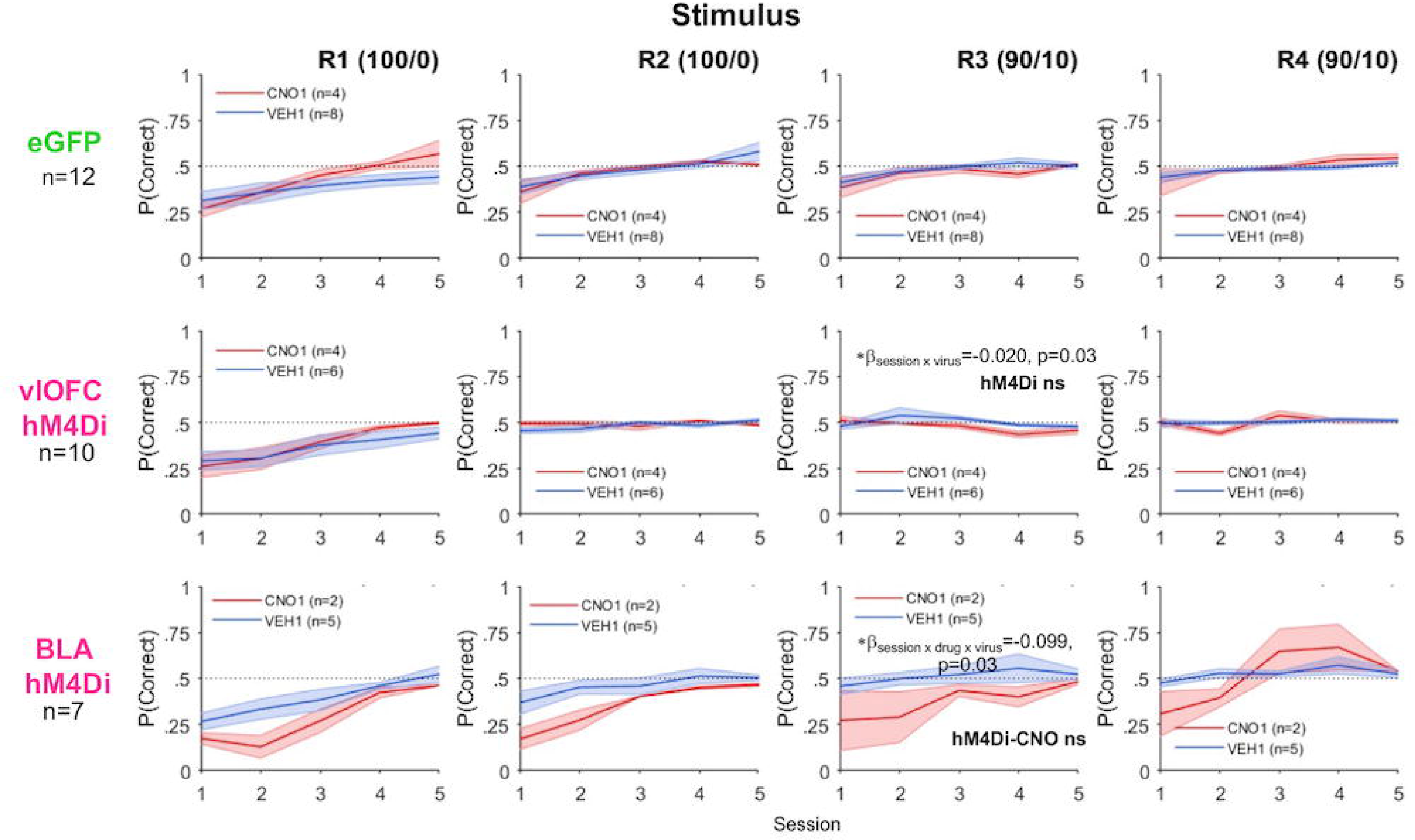
BLA inhibition further slows incremental stimulus-based reversal learning whereas vlOFC inhibition abolishes learning after first reversal. Accuracy in stimulus learners measured by mean probability correct for first 5 sessions of each deterministic (100/0) and probabilistic (90/10) reversal. Drug order was counterbalanced such that on R2 and R4 animals received VEH if they were administered CNO first on R1 and R3, and vice versa. Despite animals reaching criterion on initial learning (Fig. 2C), animals exhibited poor stimulus-based reversal learning. Reversal learning was particularly flat for R3 (first probabilistic reversal) following vlOFC inhibition compared to eGFP. Reversal learning across all reversals was especially slow for animals following BLA inhibition. Bonferroni-corrected post-hocs following a mixed-effects GLM wherein a session x virus interaction or a session x drug x virus was found resulted in *p<0.05 effect of session in eGFP, but not in hM4Di or hM4Di-CNO.

Using the above formula, for vlOFC hM4Di comparison with eGFP, we observed several interactions with virus including: session x drug x virus (β_session x drug x virus_ = −0.116, *p* = 1.64e^-03^), session x drug x drug order x virus (β_session x drug x drug order x virus_ = 0.242, *p* = 2.47e^-04^) and drug x virus x reversal number (β_drug x virus x reversal number_ = −0.133, *p* = 4.58e^-03^). With a significant interaction of session x drug x drug order x reversal number x virus (β_session x drug order x drug x virus x reversal number_ = - 0.08, *p* = 4.26e^-04^), we were justified to analyze reversals separately. We found a significant session x virus interaction only in R3 (β_session x virus_ = −0.023, *p* = 0.03). To further probe changes over session we employed the following formula for each virus separately: *γ ∼* [*1 + session + (1 + session| rat)*]. Bonferroni-corrected post-hoc comparisons revealed an effect of session in eGFP (*p* < 0.01), but not in hM4Di (*p* = 0.10), indicating that only the eGFP group improved across session in R3 (**Fig. 7**).

For BLA hM4Di compared to eGFP, fewer interactions were observed, but none which included specific reversal number. We observed an interaction of session x drug x virus (β_session x drug x virus_ = −0.098, *p* = 0.03), thus to further probe changes over session, we employed the following formula for each virus-drug combination separately: *γ ∼* [*1 + session + (1 + session| rat)*]. Bonferroni-corrected post-hoc comparisons resulted in eGFP-VEH, eGFP-CNO, and hM4Di-VEH exhibiting some improvement across sessions (session, all *p* < 0.01), but not the hM4Di-CNO group (**Fig. 7**). Thus, BLA inhibition further slowed already incremental learning across all reversals.

Due to the slow stimulus-based reversal learning, we next assessed performance measured as *probability correct* around reversals (three sessions before and after reversals for stimulus-based learning, and one session before and after reversals for action-based learning) to test for adjustments to reversals.

### Accuracy around reversals: Probability correct adjustments

*Stimulus-based reversal learning.* We analyzed accuracy (*probability correct*) around reversals to assess adjustments to reversals considering that the overall stimulus-based learning was modest. ANOVAs with virus and drug order as between subject factors were conducted on the mean change in accuracy between one reversal and the next. vlOFC hM4Di was significantly different from eGFP for R1-to-R2 [*F*_(1,18)_= 7.69, *p* = 0.01] and R2-to-R3 [*F*_(1,18)_= 7.57, *p* = 0.01], but not R3-to-R4 [*F*_(1,18)_= 1.16, *p* = 0.30], **Fig. 6D**. In contrast, BLA hM4Di was not significantly different from eGFP on changes in accuracy around any of the reversals.

*Action-based reversal learning.* For comparison, we also assessed accuracy (*probability correct*) around reversals for action-based reversal learning. As above, ANOVAs with virus and drug order as between subject factors were conducted on the mean accuracy change between one reversal and the next. Other than confirming the probabilistic reversal learning (R3) impairment for BLA hM4Di (**Fig. 2**), there were no significant effects of virus groups on accuracy changes on any other reversal transition in action-based reversal learning (**Fig. 8).**

**Figure 8.**
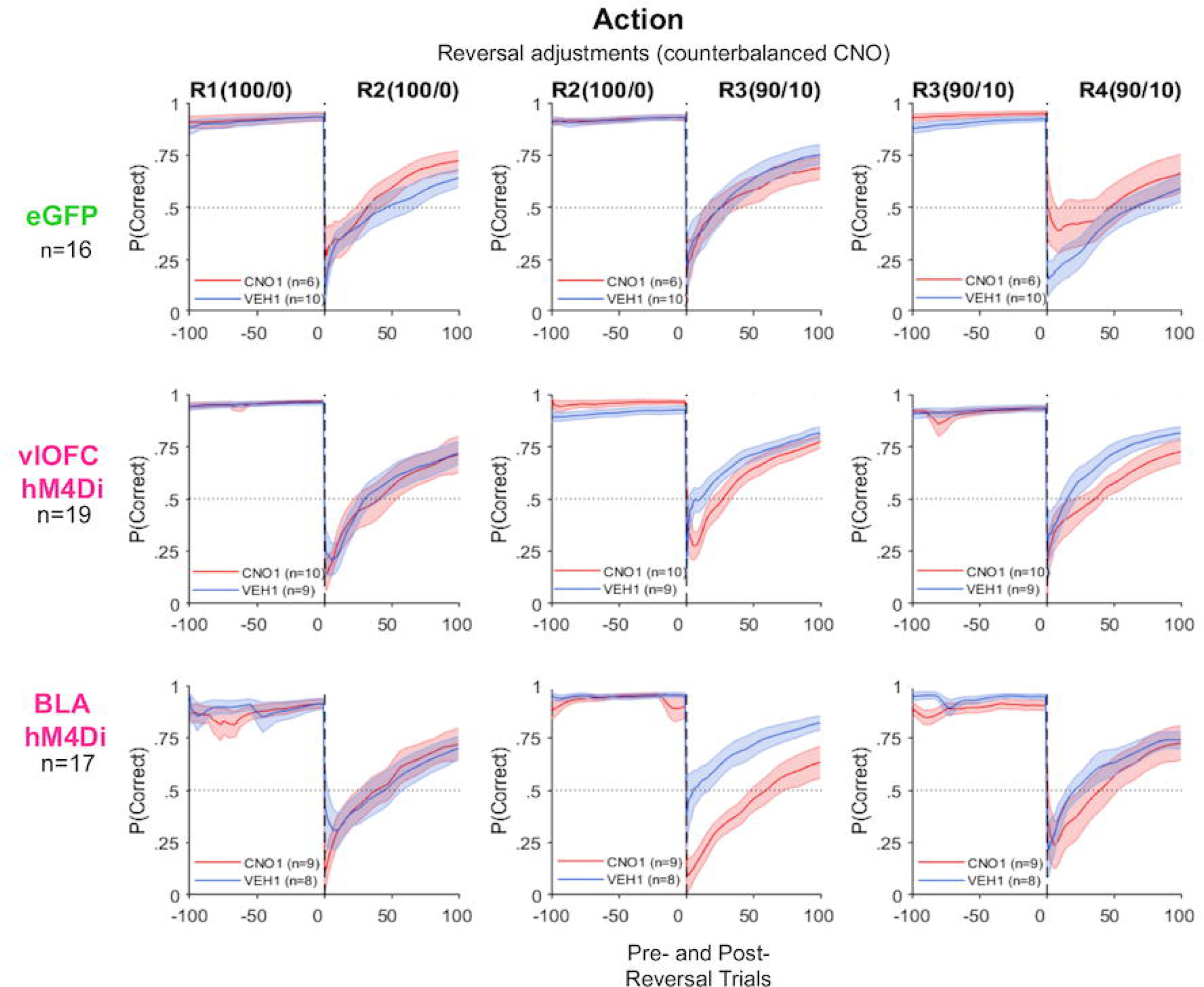
BLA, but not vlOFC, inhibition significantly slowed action-based probabilistic reversal adjustment reflected in accuracy around reversals. Plotted is the probability of choosing the better option (P(correct)) around reversal. Rats were always tested on a deterministic schedule before a probabilistic one. Shown are deterministic (100/0) and probabilistic (90/10) reversal transitions, 100 trials before and after each reversal. Drug order was counterbalanced such that on R2 and R4 animals received VEH if they were administered CNO first on R1 and R3, and vice versa. Inhibition of BLA impaired the first probabilistic reversal learning as indicated by selectively poor performance on R3 reversal. OFC inhibition resulted in no impairment around these reversals. There was also no effect of CNO in the eGFP group.

### Strategies around reversals: Win-Stay, Lose-Switch

*Stimulus-based reversal learning.* We also analyzed adaptive response strategies (*Win-Stay* and *Lose-Switch*) around reversals. ANOVAs with virus and drug order as between subject factors were conducted on mean Win-Stay or Lose-Switch between one reversal and the next. vlOFC hM4Di was significantly different from eGFP for *Win-Stay* R1-to-R2 [*F*_(1,18)_ =8.97, *p* < 0.01] and R2-to-R3 [*F*_(1,18)_= 7.81, *p* = 0.01], but not R3-to-R4 [*F*_(1,18)_= 0.40, *p* = 0.54]. However, though there was a trend for reduced Lose-Switch in vlOFC hM4Di for R1-to-R2 [*F*_(1,18)_= 3.64, *p* = 0.07] and R2-to-R3 [*F*_(1,18)_= 3.69, *p* = 0.07], this group did not statistically differ from eGFP on this measure, **Fig. 9**.

**Figure 9.**
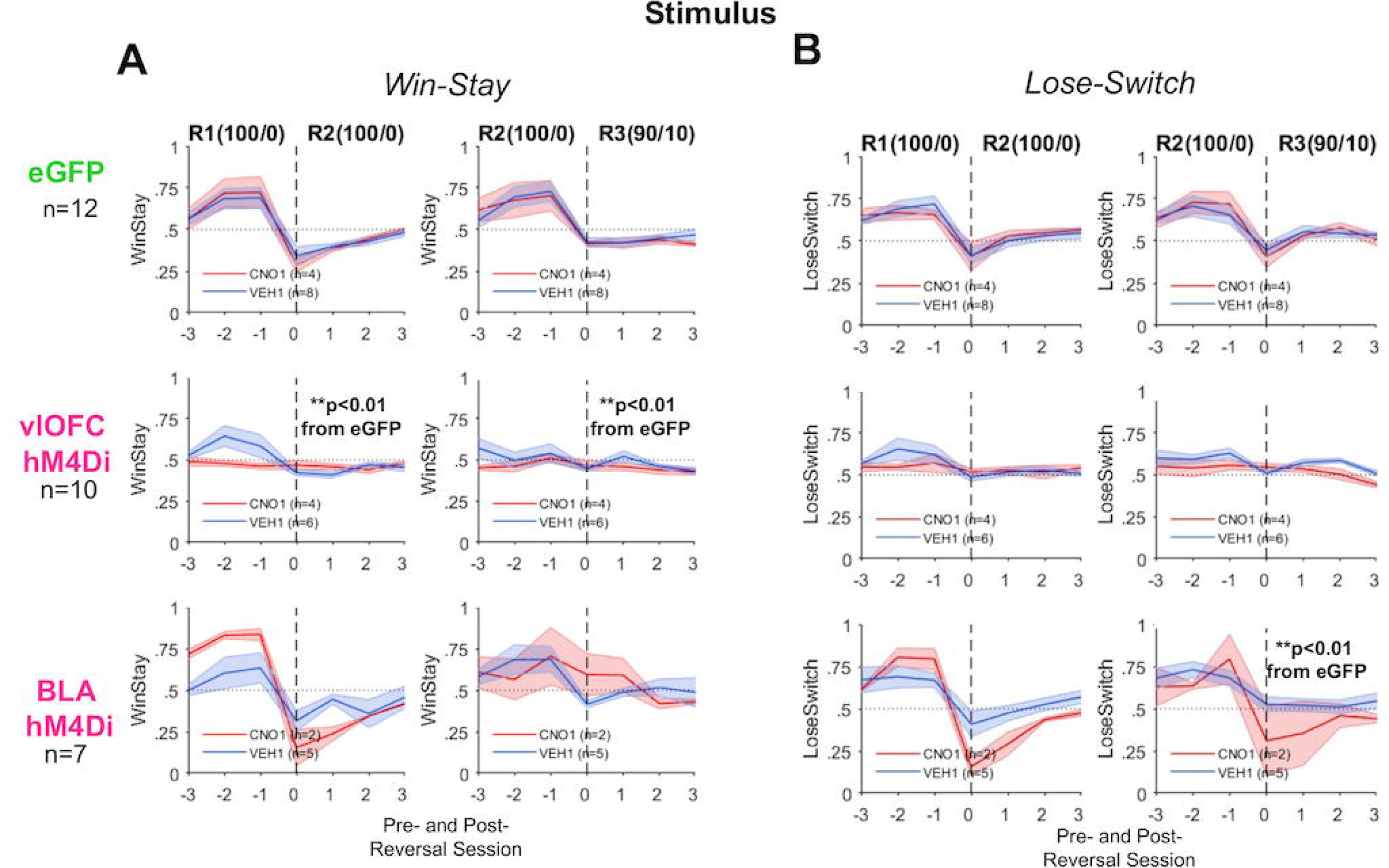
vlOFC, but not BLA, inhibition impairs adjustments to deterministic stimulus-based reversals as measured by changes in Win-Stay strategies. Plots of Win-Stay and Lose-Switch around reversals were restricted to include only animals that reached greater than 50% running window average for the last 100 trials in initial discrimination. **(A)** Inhibition of vlOFC abolishes changes in Win-Stay over the last 3 (pre) and first 3 (post)-reversal sessions, indicating impaired R1-R2 and R2-R3 transitions. In contrast, BLA inhibition had no impact on performance during these transitions. **(B)** Inhibition of BLA attenuates changes in Lose-Switch over the last 3 (pre) and first 3 (post)-reversal sessions in R2-R3, or the transition from deterministic to probabilistic schedule. There was also no effect of CNO in eGFP group learning. **p<0.01 different than eGFP following ANOVA of pre-post difference.

In contrast, BLA hM4Di was significantly different from eGFP on changes in *Lose-Switch* strategies around only R2-to-R3 [*F*_(1,15)_= 5.82, *p* = 0.03]. The results for these adaptive strategies reflect a similar pattern to that observed for *probability correct* for both vlOFC and BLA hM4Di, above.

In summary, our results indicate that BLA, but not vlOFC, is required for learning probabilistic reversals. Conversely, vlOFC, but not BLA, is necessary for rapid adjustments to reversals, generally.

## Discussion

We used a chemogenetic approach to transiently inactivate neurons in either vlOFC or BLA to assess how these regions are involved in different aspects of reversal learning. Although the role of OFC in reversal learning has been instantiated in different paradigms using visual stimuli and cues (Izquierdo et al., 2013; Piantadosi et al., 2018; Hervig et al., 2020; Alsio et al., 2021) as well as olfactory ones (Schoenbaum et al., 2003; Kim and Ragozzino, 2005), several groups also report a strong role for OFC in action (spatial)-based reversal learning (Dalton et al., 2016; Groman et al., 2019; Verharen et al., 2020). Almost all of these reversal learning investigations have involved irreversible lesions or baclofen/muscimol inactivations of OFC. Testing both types with a chemogenetic approach targeting projection neurons, here we found that rapid adjustments to reversals generally rely on vlOFC, across both stimuli and actions.

In parallel, the specific role of BLA in stimulus-vs. action-based reversal learning is poorly understood given mixed results (Schoenbaum et al., 2003; Izquierdo and Murray, 2004; Churchwell et al., 2009; Hervig et al., 2020). Recent studies suggest amygdala may be involved in both types of learning (Taswell et al., 2021; Keefer and Petrovich, 2022) as BLA activity is modulated by violations in reward expectations generally, which are not association-specific (Roesch et al., 2012). To probe this, we tested animals on both stimulus- and action-based tasks and found that BLA is required for both stimulus- and action-based reversal learning, with a more pronounced role in probabilistic reversal learning (i.e., detecting meaningful change against a background of uncertainty). This adds to some empirical evidence (Stolyarova & Izquierdo, 2017) and supports its theorized role in learning under unexpected uncertainty (Soltani & Izquierdo, 2019).

As additional motivations for the present study, several reports suggest that neural recruitment in reversal learning may depend on certainty of rewards (Boulougouris et al., 2007; Boulougouris and Robbins, 2009; Ward et al., 2015; Costa et al., 2016; Dalton et al., 2016; Piantadosi et al., 2018; Verharen et al., 2020). To further understand this, we tested animals on both deterministic (100/0) and probabilistic reversals (90/10). We found vlOFC to be involved in the rapid adjustment to stimulus-based reversals and in the initial learning of both deterministic and probabilistic learning of actions, whereas BLA was more selectively involved in probabilistic reversal learning, not adjustments to reversals.

Finally, due to the sparsity of research probing sex differences in flexible learning and decision making (Orsini and Setlow, 2017; Grissom and Reyes, 2019; Orsini et al., 2022; Cox et al., 2023), where an overwhelming number of reversal learning studies include only males (Schoenbaum et al., 2003; Izquierdo et al., 2013; Dalton et al., 2016; Groman et al., 2019; Hervig et al., 2020; Verharen et al., 2020), we included both male and female rats here. We found sex-dependent contributions of vlOFC in early action-based reversal learning. We elaborate on these findings within the context of the existing literature below.

### Preferential recruitment of BLA over vlOFC during probabilistic reversal learning

All animals learned to flexibly adjust their responses following deterministic and probabilistic action-based reversals, indicating successful remapping of reward contingencies as accuracy increased across trials. Importantly, we found no effect of CNO in eGFP animals, suggesting that it was activation of hM4Di receptors in BLA that were crucial to any impairments observed. vlOFC was especially involved in early learning of first deterministic (R1) and probabilistic reversal (R3). This is consistent with findings following pharmacological inactivations or lesions of OFC (Boulougouris et al., 2007; Boulougouris and Robbins, 2009; Dalton et al., 2016; Piantadosi et al., 2018; Verharen et al., 2020).

BLA inhibition was not expected to impair deterministic reversal learning as it is thought to be mostly recruited when there is some level of uncertainty, e.g., probabilistic outcomes (Roesch et al., 2012), and we found evidence of that here. Amygdala-lesioned monkeys are also impaired on action(spatial)-based probabilistic reversal learning, exhibiting decreased probability of choosing the better option, and increased switching behavior following negative outcomes (Taswell et al., 2021). BLA may indeed be critical in generating prediction error signals following changes in reward associations (Esber et al., 2012; Roesch et al., 2012; Iordanova et al., 2021), with particular involvement in detecting unexpected upshifts or downshifts in value (Roesch et al., 2010; Stolyarova and Izquierdo, 2017). Our finding of attenuated learning of probabilistic reversal R3 suggests it is the reversal superimposed on the misleading feedback that most engages BLA. Here we also probed if BLA was necessary during the initial learning of probabilistic outcomes, without reversal experience (**Fig. 4**). Inhibition during initial probabilistic learning did not produce an impairment, indicating BLA is required during the convergence of both probabilistic feedback and shift in contingency (i.e., unexpected uncertainty).

### vlOFC, but not BLA, is necessary for adjustments to reversals

As described above, unlike the ease of action-based reversal learning, rats exhibited difficulty learning reversals of stimulus-reward contingencies, as previously reported (Harris, Aguirre et al., 2021). Thus, instead of examining acquisition curves which reached asymptote slightly above chance, we elected to study adjustment to reversals by comparing accuracy and strategy prior to and after a reversal occurred. Furthermore, this enabled assessment about whether prior inhibition affected future adjustments to reversals and whether this varied by transition type [i.e., between deterministic reversals (R1→ R2), deterministic and probabilistic reversal (R2→R3), or probabilistic reversals (R3→R4)]. We found that vlOFC, but not BLA, inhibition produced a failure in detecting first deterministic and first probabilistic reversal. This pattern was not observed in animals that received VEH during the first deterministic reversal, suggesting vlOFC needs to be “online” when experiencing a reversal for the very first time as this determines how flexibly animals respond to future reversals. Employment of adaptive strategies matched this effect, such that vlOFC-inhibited animals did not employ effective win-stay strategies after the first reversal. That vlOFC inhibition did not impair the ability to adjust to probabilistic reversals (R3→R4) supports the idea that other brain regions may be recruited when the probabilistic reward contingencies have already been established. The role of OFC in establishing an “expected uncertainty” (Soltani and Izquierdo, 2019) has been instantiated experimentally in several recent studies using different methodologies (Stolyarova & Izquierdo, 2017; Namboodiri et al., 2019; Namboodiri et al., 2021; Jenni et al., 2022), and we add the establishment of expected uncertainty in adjustments to stimulus-based reversals to this evidence.

In contrast, BLA inhibition did not result in any impairment in the ability to adjust to reversals, with the exception of the transition to a probabilistic schedule (i.e., in lose-switch strategies), which lends additional support to the idea that BLA is particularly engaged in adjusting behavior to meaningful changes in the reward environment. Our results based on estimated RL model parameters suggests differential mechanisms for adjustment to reversals between males and females and following vlOFC vs. BLA inhibition. Specifically, we found evidence for differential effects on γ_d_, where the decay rate for unchosen action values was greater for females than males following vlOFC inhibition. This is consistent with a previous study which also reported similar disruption in retention of action values after ablating OFC neurons projecting to BLA (Groman et al., 2019). Notably, Groman et al. (2019) included only male rats and involved a stronger manipulation than our chemogenetic approach (i.e., one that caused pathway-specific neuronal apoptosis). Together with our findings, we can conclude that neurons in OFC, but not BLA, store a memory of action values that are used to adjust to reversals. These results stand in contrast to the maladaptive, increased choice consistency following BLA inhibition that instead reflects poor updating.

### Stimulus-based vs. action-based learning

Interestingly, we discovered task order to be significant in rats’ ability to learn to discriminate stimuli: stimulus→action was learned much more readily than action→stimulus. This can be explained by noting that rats are heavily biased to acquire spatial associations (Wright et al., 2019), and reinforcing this already-strong learning likely inhibits the ability to learn associations where spatial information should be ignored. In contrast, nonhuman primates are able to quickly transition between “what” vs. “where” blocks of trials (Rothenhoefer et al., 2017; Taswell et al., 2021). Nonetheless, learning both types of associations are crucial for flexibility required in naturalistic environments and thus, it is important to examine how stimulus-based and action-based learning systems interact with each other (Soltani & Koechlin, 2022). Moreover, although the role of OFC in stimulus-or cue-based reversal learning has been probed using olfactory and visual stimuli, more viral-mediated approaches employing targeted chemogenetic and optogenetic manipulations across sensory modalities in both males and females are warranted.

## Conclusion

The present results indicate dissociable roles for vlOFC and BLA in flexible learning under uncertainty. BLA is crucial in probabilistic reversal learning, or learning of unexpected changes against a background uncertainty, whereas vlOFC is important in the establishment of expected uncertainty that enables adjustments to reversals. Interestingly, females exhibited a larger memory decay for the unchosen option following vlOFC inhibition than males, indicating a sex-dependent influence on learning under uncertainty. Future investigations could combine targeted causal manipulations with neural correlate approaches to assess the timescales and dynamics of cortico-amygdalar involvement in this learning.

## Acknowledgements

This work was supported by UCLA’s Division of Life Sciences Retention fund (Izquierdo), National Institutes of Health grants R01 DA047870 (Izquierdo and Soltani), R21 MH122800 (Izquierdo and Blair), R01AA024527 (Spigelman), K01 DA042219 (O’Neill), and F31 AA028183 (Munier), the NSF GRFP, Cota-Robles Fellowship, and Charles E. and Sue K. Young Fellowship (Aguirre), Ursula Mandel Fellowship and Graduate Research Mentorship award (Romero-Sosa), and the Training program in Neurotechnology Translation T32 NS115753 (Ye). We acknowledge the Staglin Center for Brain and Behavioral Health for additional support related to fluorescence microscopy. We thank P. Ganupuru for assistance with brain collection. We also thank the NIDA Drug Supply program for the supply of clozapine-N-oxide.

## Code availability

Code and data will be made available upon publication https://github.com/izquierdolab and https://gin.g-node.org/aizquie

**The authors declare no competing financial interests.**

## Notes

### Competing Interest Statement

The authors have declared no competing interest.

### Summary of Updates

Additional control experiments and corresponding figures included

## References

Alsio J, Lehmann O, McKenzie C, Theobald DE, Searle L, Xia J, Dalley JW, Robbins TW (2021) Serotonergic Innervations of the Orbitofrontal and Medial-prefrontal Cortices are Differentially Involved in Visual Discrimination and Reversal Learning in Rats. Cereb Cortex 31:1090–1105.

Amodeo LR, McMurray MS, Roitman JD (2017). Orbitofrontal cortex reflects changes in response-outcome contingencies during probabilistic reversal learning. Neuroscience 345:27–37.

Asrican B, Song J (2021) Extracting meaningful circuit-based calcium dynamics in astrocytes and neurons from adult mouse brain slices using single-photon GCaMP imaging. STAR Protoc 2:100306.

Barreiros IV, Panayi MC, Walton ME (2021a) Organization of Afferents along the Anterior-posterior and Medial-lateral Axes of the Rat Orbitofrontal Cortex. Neuroscience 460:53–68.

Barreiros IV, Ishii H, Walton ME, Panayi MC (2021b) Defining an orbitofrontal compass: Functional and anatomical heterogeneity across anterior-posterior and medial-lateral axes. Behav Neurosci 135:165–173.

Behrens TE, Woolrich MW, Walton ME, Rushworth MF (2007) Learning the value of information in an uncertain world. Nat Neurosci 10:1214–1221.

Boulougouris V, Robbins TW (2009) Pre-surgical training ameliorates orbitofrontal-mediated impairments in spatial reversal learning. Behavioural Brain Research 197:469–475.

Boulougouris V, Dalley JW, Robbins TW (2007) Effects of orbitofrontal, infralimbic and prelimbic cortical lesions on serial spatial reversal learning in the rat. Behavioural Brain Research 179:219–228.

Churchwell JC, Morris AM, Heurtelou NM, Kesner RP (2009) Interactions between the prefrontal cortex and amygdala during delay discounting and reversal. Behav Neurosci 123:1185–1196.

Corbit LH, Balleine BW (2005) Double dissociation of basolateral and central amygdala lesions on the general and outcome-specific forms of pavlovian-instrumental transfer. J Neurosci 25:962–970.

Costa KM, Scholz R, Lloyd K, Moreno-Castilla P, Gardner MPH, Dayan P, Schoenbaum G (2023) The role of the lateral orbitofrontal cortex in creating cognitive maps. Nat Neurosci 26:107–115.

Costa VD, Dal Monte O, Lucas DR, Murray EA, Averbeck BB (2016) Amygdala and Ventral Striatum Make Distinct Contributions to Reinforcement Learning. Neuron 92:505–517.

Cox J, Minerva AR, Fleming WT, Zimmerman CA, Hayes C, Zorowitz S, Bandi A, Ornelas S, McMannon B, Parker NF, Witten IB (2023) A neural substrate of sex-dependent modulation of motivation. Nat Neurosci 26(2):274–284.

Dalton GL, Wang NY, Phillips AG, Floresco SB (2016) Multifaceted Contributions by Different Regions of the Orbitofrontal and Medial Prefrontal Cortex to Probabilistic Reversal Learning. J Neurosci 36:1996–2006.

Esber GR, Roesch MR, Bali S, Trageser J, Bissonette GB, Puche AC, Holland PC, Schoenbaum G (2012) Attention-related Pearce-Kaye-Hall signals in basolateral amygdala require the midbrain dopaminergic system. Biol Psychiatry 72:1012–1019.

Ghasemi A, Jeddi S, Kashfi K (2021) The laboratory rat: Age and body weight matter. EXCLI J 20:1431–1445.

Grissom NM, Reyes TM (2019) Let’s call the whole thing off: evaluating gender and sex differences in executive function. Neuropsychopharmacology 44:86–96.

Groman SM, Keistler C, Keip AJ, Hammarlund E, DiLeone RJ, Pittenger C, Lee D, Taylor JR (2019) Orbitofrontal Circuits Control Multiple Reinforcement-Learning Processes. Neuron 103:734–746 e733.

Harris C, Aguirre C, Kolli S, Das K, Izquierdo A, Soltani A (2021) Unique features of stimulus-based probabilistic reversal learning. Behav Neurosci 135:550–570.

Hart EE, Blair GJ, O’Dell TJ, Blair HT, Izquierdo A (2020) Chemogenetic Modulation and Single-Photon Calcium Imaging in Anterior Cingulate Cortex Reveal a Mechanism for Effort-Based Decisions. J Neurosci 40:5628–5643.

Hervig ME, Fiddian L, Piilgaard L, Bozic T, Blanco-Pozo M, Knudsen C, Olesen SF, Alsio J, Robbins TW (2020) Dissociable and Paradoxical Roles of Rat Medial and Lateral Orbitofrontal Cortex in Visual Serial Reversal Learning. Cereb Cortex 30:1016–1029.

Iordanova MD, Yau JO-Y, McDannald MA, Corbit LH (2021) Neural substrates of appetitive and aversive prediction error. Neurosci Biobehav Rev 123:337–351.

Izquierdo A (2017) Functional Heterogeneity within Rat Orbitofrontal Cortex in Reward Learning and Decision Making. J Neurosci 37:10529–10540.

Izquierdo A, Murray EA (2004) Combined Unilateral Lesions of the Amygdala and Orbital Prefrontal Cortex Impair Affective Processing in Rhesus Monkeys. Journal of Neurophysiology 91:2023–2039.

Izquierdo A, Darling C, Manos N, Pozos H, Kim C, Ostrander S, Cazares V, Stepp H, Rudebeck PH (2013) Basolateral amygdala lesions facilitate reward choices after negative feedback in rats. J Neurosci 33:4105–4109.

Janak PH, Tye KM (2015) From circuits to behaviour in the amygdala. Nature 517:284–292.

Jang AI, Costa VD, Rudebeck PH, Chudasama Y, Murray EA, Averbeck BB (2015) The Role of Frontal Cortical and Medial-Temporal Lobe Brain Areas in Learning a Bayesian Prior Belief on Reversals. J Neurosci 35:11751-11760.

Jenni NL, Rutledge G, Floresco SB (2022) Distinct Medial Orbitofrontal-Striatal Circuits Support Dissociable Component Processes of Risk/Reward Decision-Making. J Neurosci 42:2743–2755.

Jones B, Mishkin M (1972). Limbic lesions and the problem of stimulus-reinforcement associations. Experimental Neurology 36:362–377.

Keefer SE, Petrovich GD (2022) Necessity and recruitment of cue-specific neuronal ensembles within the basolateral amygdala during appetitive reversal learning. Neurobiol Learn Mem 194:107663.

Kim J, Ragozzino ME (2005) The involvement of the orbitofrontal cortex in learning under changing task contingencies. Neurobiol Learn Mem 83:125–133.

Lichtenberg NT, Pennington ZT, Holley SM, Greenfield VY, Cepeda C, Levine MS, Wassum KM (2017) Basolateral Amygdala to Orbitofrontal Cortex Projections Enable Cue-Triggered Reward Expectations. J Neurosci 37:8374–8384.

Malvaez M, Shieh C, Murphy MD, Greenfield VY, Wassum KM (2019) Distinct cortical-amygdala projections drive reward value encoding and retrieval. Nat Neurosci 22:762–769.

Namboodiri VMK, Hobbs T, Trujillo-Pisanty I, Simon RC, Gray MM, Stuber GD (2021) Relative salience signaling within a thalamo-orbitofrontal circuit governs learning rate. Curr Biol 31:5176–5191.e5175.

Namboodiri VMK, Otis JM, van Heeswijk K, Voets ES, Alghorazi RA, Rodriguez-Romaguera J, Mihalas S, Stuber GD (2019) Single-cell activity tracking reveals that orbitofrontal neurons acquire and maintain a long-term memory to guide behavioral adaptation. Nat Neurosci 22:1110–1121.

Orsini CA, Setlow B (2017) Sex differences in animal models of decision making. J Neurosci Res 95:260–269.

Orsini CA, Truckenbrod LM, Wheeler A-R (2022) Regulation of sex differences in risk-based decision making by gonadal hormones: Insights from rodent models. Behav Processes 200:104663–104663.

Pachitariu M, Stringer C, Dipoppa M, Schröder S, Rossi LF, Dalgleish H, Carandini M, Harris KD (2017) Suite2p: beyond 10,000 neurons with standard two-photon microscopy. bioRxiv:061507.

Piantadosi PT, Lieberman AG, Pickens CL, Bergstrom HC, Holmes A (2018) A novel multichoice touchscreen paradigm for assessing cognitive flexibility in mice. Learn Mem 26:24–30.

Roesch MR, Calu DJ, Esber GR, Schoenbaum G (2010) Neural correlates of variations in event processing during learning in basolateral amygdala. J Neurosci 30:2464–2471.

Roesch MR, Esber GR, Bryden DW, Cerri DH, Haney ZR, Schoenbaum G (2012) Normal aging alters learning and attention-related teaching signals in basolateral amygdala. J Neurosci 32:13137–13144.

Rothenhoefer KM, Costa VD, Bartolo R, Vicario-Feliciano R, Murray EA, Averbeck BB (2017) Effects of ventral striatum lesions on stimulus-based versus action-based reinforcement learning. J Neurosci 37: 6902–6914.

Rudebeck PH, Murray EA (2008) Amygdala and orbitofrontal cortex lesions differentially influence choices during object reversal learning. J Neurosci 28:8338–8343.

Schoenbaum G, Saddoris MP, Stalnaker TA (2007). Reconciling the roles of orbitofrontal cortex in reversal learning and the encoding of outcome expectancies. Ann N Y Acad Sci 1121:320–35.

Schoenbaum G, Setlow B, Nugent SL, Saddoris MP, Gallagher M (2003) Lesions of orbitofrontal cortex and basolateral amygdala complex disrupt acquisition of odor-guided discriminations and reversals. Learn Mem 10:129–140.

Sias AC, Morse AK, Wang S, Greenfield VY, Goodpaster CM, Wrenn TM, Wikenheiser AM, Holley SM, Cepeda C, Levine MS, Wassum KM (2021) A bidirectional corticoamygdala circuit for the encoding and retrieval of detailed reward memories. Elife 10.

Soltani A, Izquierdo A (2019) Adaptive learning under expected and unexpected uncertainty. Nat Rev Neurosci 20:635–644.

Soltani A, Koechlin E (2022) Computational models of adaptive behavior and prefrontal cortex. Neuropsychopharmacology 47: 58–71.

Stalnaker TA, Roesch MR, Calu DJ, Burke KA, Singh T, Schoenbaum G (2007) Neural correlates of inflexible behavior in the orbitofrontal-amygdalar circuit after cocaine exposure. Ann N Y Acad Sci 1121:598–609.

Stolyarova A, Izquierdo A (2017) Complementary contributions of basolateral amygdala and orbitofrontal cortex to value learning under uncertainty. Elife 6.

Stolyarova A, Rakhshan M, Hart EE, O’Dell TJ, Peters MAK, Lau H, Soltani A, Izquierdo A (2019) Contributions of anterior cingulate cortex and basolateral amygdala to decision confidence and learning under uncertainty. Nat Commun 10:4704.

Tada M, Takeuchi A, Hashizume M, Kitamura K, Kano M (2014) A highly sensitive fluorescent indicator dye for calcium imaging of neural activity in vitro and in vivo. Eur J Neurosci 39:1720–1728.

Taswell CA, Costa VD, Basile BM, Pujara MS, Jones B, Manem N, Murray EA, Averbeck BB (2021) Effects of Amygdala Lesions on Object-Based Versus Action-Based Learning in Macaques. Cereb Cortex 31:529–546.

Tye KM, Janak PH (2007) Amygdala neurons differentially encode motivation and reinforcement. J Neurosci 27:3937–3945.

Verharen JPH, den Ouden HEM, Adan RAH, Vanderschuren L (2020) Modulation of value-based decision making behavior by subregions of the rat prefrontal cortex. Psychopharmacology (Berl) 237:1267–1280.

Ward RD, Winiger V, Kandel ER, Balsam PD, Simpson EH (2015) Orbitofrontal cortex mediates the differential impact of signaled-reward probability on discrimination accuracy. Front Neurosci 9:230.

Wassum KM, Izquierdo A (2015) The basolateral amygdala in reward learning and addiction. Neurosci Biobehav Rev 57:271–283.

Winstanley CA, Floresco SB (2016) Deciphering Decision Making: Variation in Animal Models of Effort-and Uncertainty-Based Choice Reveals Distinct Neural Circuitries Underlying Core Cognitive Processes. J Neurosci 36:12069–12079.

Wright SL, Martin GM, Thorpe CM, Haley K, Skinner DM (2019) Distance and direction, but not light cues, support response reversal learning. Learning & Behavior 47:38–46.

Ye T, Romero-Sosa JL, Rickard A, Aguirre CG, Wikenheiser AM, Blair HT, Izquierdo A (2023) Theta oscillations in anterior cingulate cortex and orbitofrontal cortex differentially modulate accuracy and speed in flexible reward learning. Oxford Open Neuroscience kvad005.

Zimmermann KS, Li CC, Rainnie DG, Ressler KJ, Gourley SL (2018) Memory Retention Involves the Ventrolateral Orbitofrontal Cortex: Comparison with the Basolateral Amygdala. Neuropsychopharmacology 43:373–383.

